# HIF1A recruits primate-specific endogenous retroviruses into the human hypoxic and immune responses

**DOI:** 10.64898/2026.08.31.748411

**Authors:** Umut Cakir, Federica Mantovani, Souad Youjil Abadi, Joachim Fandrey, Sandra Winning, Hannelore Ehrenreich, Hannah S. Schwarzer-Sperber, Zsuzsanna Izsvak, Roland Schwarzer, Manvendra Singh

**Author notes:** Corresponding authors Prof. Dr. Zsuzsanna Izsvák, Mobile DNA Group, Max-Delbrück-Center for Molecular Medicine, Robert-Rössle-Str. 10, 13125 Berlin-Buch, Berlin, Germany, Dr. Roland Schwarzer, Institute for the Research on HIV and AIDS-Associated Diseases (HIV-AAD), University Hospital Essen, University Duisburg-Essen, Essen, Germany, Dr. Manvendra Singh, PhD, Institut Necker Enfants Malades (INEM), Université Paris Cité, INSERM UMR 1151, Paris 75015, France.

## Abstract

Oxygen availability varies profoundly across the human body and changes further during inflammation, infection, tissue injury and disease. Immune cells must therefore continuously adapt their transcriptional and metabolic state based on the oxygen availability to them. Hypoxia-inducible factor 1α (HIF1A) is central to this adaptation and a marker of the cellular response to low oxygen, yet its genomic targets have been assembled from a non-repetitive fraction of the genome, leaving nearly half of the human genome largely unexplored. Here we define the gene and transposable-element (TE) landscape of the human hypoxic response across different human tissues, cell lines, and conditions. This directional TE response was reproduced in transformed cells and in primary immune cells isolated from blood and the physiologically oxygen-restricted tonsil. Single-cell profiling of peripheral blood mononuclear cells (PBMC) under hypoxia, pharmacological HIF stabilization, and interferon stimulation revealed a striking difference between the gene and retrotranscriptome responses. While gene responses were strongly cell-type dependent and in a bidirectional manner, TEs were overwhelmingly activated. This pattern extended to blood and tonsil immune cells, where ∼70-90% of tested TE families were induced under hypoxia, with activated tonsil cells showing exclusively induced significant families, including THE1B, alongside increased LTR7 and HERVH.

Integrating HIF1A ChIP-seq with transcriptional responses revealed that HIF1A does not engage repetitive DNA indiscriminately. Instead, its binding converged on LTR7, the promoter long terminal repeat of the HERVH endogenous retrovirus. Approximately 80% of HIF1A-bound LTR7 elements contained a canonical hypoxia-response element, and disruption of HIF1A DNA binding dramatically reduced the expression of occupied HERVH loci. CRISPR deletion of individual LTR7/HERVH loci altered the expression of distant and neighboring genes, demonstrating that hypoxia-responsive retroelements can participate directly in host gene regulation and contribute to overall physiology. Our findings reveal the repetitive genome as a previously underappreciated component of oxygen sensing. We propose that HIF1A recruits selected endogenous retroviral elements into the human hypoxic response, extending oxygen-dependent regulation beyond conventional gene promoters and providing an additional regulatory layer through which tissue oxygenation can shape immune-cell state and human physiology.

## INTRODUCTION

Oxygen is not uniformly distributed throughout the human body. Its availability differs substantially between organs, within individual tissues, and across physiological states, creating a local environment in which cells must continuously adapt^1^. It even varies during physical activity, inflammation, infection, tissue injury, and tumour growth^2,3^. This is particularly relevant to the immune system. Immune cells originate, circulate, and function across markedly different oxygen environments, from relatively oxygen-rich blood to the lower oxygen tensions of bone marrow, lymphoid tissues, and inflamed lesions. Oxygen sensing is consequently intertwined with immune-cell metabolism, differentiation, and effector function^4–6^.

When oxygen becomes limiting, cells use a conserved transcriptional program regulated by hypoxia-inducible factors (HIFs)^7–9^. Under normoxia, the HIF1A subunit undergoes continuous post-translational hydroxylation at proline residues by prolyl hydroxylase domain enzymes and subsequently degrades^10–12^. As oxygen decreases, hydroxylation is reduced, and HIF1A proteins accumulate, and they enter the nucleus. In the nucleus, it heterodimerizes with HIF-1β/ARNT^13,14^, and binds hypoxia response elements (HREs) to regulate the genes controlling glycolytic reprogramming, angiogenesis, cell survival, and numerous other adaptive processes (Figure 1A)^15,16^. This pathway is one of the most fundamental mechanisms by which mammalian cells sense their environment. The HRE, however, is not a simple on/off switch. HRE sequence composition, local chromatin accessibility, and cooperating transcription factors, including AP-1 and NF-κB, influence where HIF1A binds and which genes respond^17–20^. Thus, the hypoxic response is not a fixed transcriptional programme but is shaped by cellular identity and tissue context.

**Figure 1.**
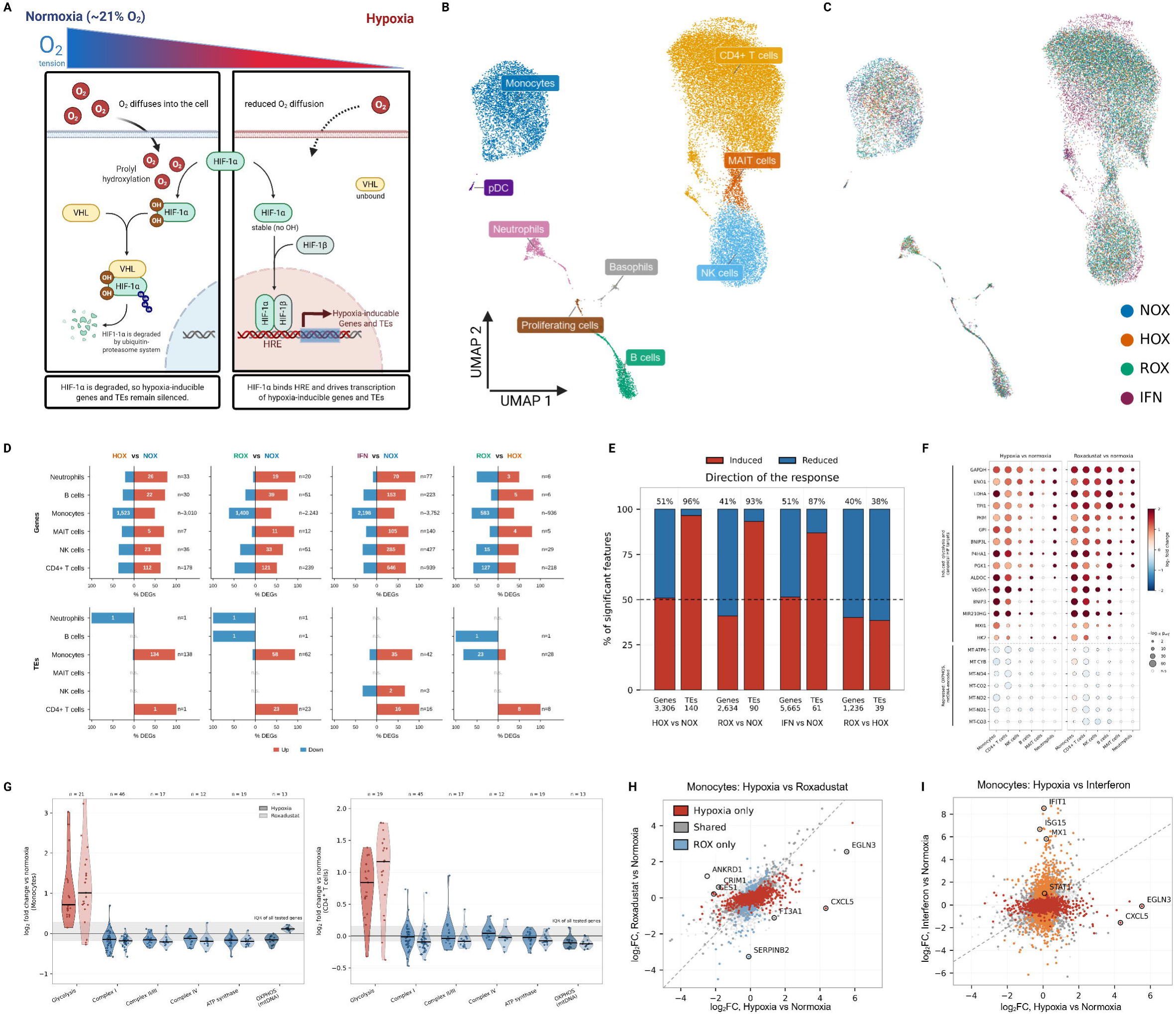
Single-cell profiling separates a cell-type-specific gene response from a uniformly activating TE response. **(A)** The HIF1A is a master transcription factor in the oxygen-sensing pathway. Under normoxia, HIF-1A is prolyl-hydroxylated and degraded through the VHL ubiquitin-proteasome pathway, so hypoxia-inducible genes and TEs remain silent. Under hypoxia, HIF-1A escapes hydroxylation, dimerises with HIF-1β (ARNT), and binds hypoxia response elements (HREs). **(B)** UMAP of PBMCs from three donors profiled by single-cell transcriptome and AbSeq protein sequencing, coloured by annotated cell type. Nine populations were annotated. These are CD4⁺ T cells, NK cells, monocytes, B cells, MAIT cells, neutrophils, basophils, plasmacytoid dendritic cells and a proliferating population. **(C)** The same cells show on (B) coloured by condition: normoxia (NOX), hypoxia (HOX, 1% O₂), roxadustat (ROX) and interferon (IFN). **(D)** Number and direction of significantly differentially expressed genes (top) and TE families (bottom) for each of four comparisons (padj < 0.05), shown for the six populations with sufficient cells for testing: monocytes, CD4⁺ T cells, NK cells, B cells, MAIT cells and neutrophils. Bars show the percentage induced and reduced; n gives the total per one cell type. **(E)** Direction of the response, expressed as the percentage of significant features induced, for genes and TE families in each comparison. Percentages above each bar give the induced proportions. (F) Expression of induced glycolytic and canonical HIF target genes (top) and repressed mitochondrially encoded OXPHOS genes (bottom) across the same six populations under hypoxia and roxadustat. Point size reflects −log₁₀ padj and colour the log₂ fold changes. **(G)** Pathway-level shifts in monocytes (left) and CD4⁺ T cells (right). Each violin shows the distribution of log₂ fold changes for all tested genes in that pathway relative to normoxia, with the interquartile range of these genes shaded for reference. Colour denotes the pathway red for glycolysis and blue for the five oxidative phosphorylation sets. n gives the number of genes per set. **(H)** Monocyte gene-level comparison of hypoxia and roxadustat. Points are coloured by whether they reach significance under hypoxia only, roxadustat only, or both. Selected genes separating the two stimuli are labelled. **(I)** Monocyte gene-level comparison of hypoxia and interferon, coloured in the same way as in (H).

This context dependency is particularly evident in immunity. Physiologically, oxygen-restricted niches occur in bone marrow, intestinal mucosa, and tonsillar germinal centres, while inflammation can generate even more severe local hypoxia. Furthermore, circulating immune cells must rapidly adapt to sharp drops in oxygen as they migrate from well-oxygenated blood into oxygen-starved tissues. Thus, hypoxia is a routine rather than an exceptional feature of the environment in which immune cells operate^5^. Effector memory T cells, for instance, proliferate and kill target cells more efficiently under hypoxia in association with their glycolytic state and HIF1A activity^21^. HIF1A, therefore, acts as a physiological regulator of immune gene expression under conditions that immune cells routinely encounter.

Despite decades of work defining HIF biology, an important part of the HIF-regulated genome remains largely unexplored. Nearly half of the human genome derives from transposable elements (TEs), including LINEs, SINEs, and long terminal repeat (LTR) elements derived from endogenous retroviruses (ERVs)^22,23^. Once dismissed as genomic parasites, TEs are now recognized as a major source of regulatory innovations^24,25^. LTR elements, in particular, are pre-equipped with transcription factor (TF) binding sites and can function as promoters, enhancers, insulators and alternative transcriptional units, allowing evolutionarily related groups of elements to distribute regulatory sequences across the genome. This property has been particularly important in the evolution of innate immunity, where ERV-derived elements such as MER41 have provided interferon-responsive enhancers for host defence genes. Retroelements, therefore, offer a mechanism through which ancient viral sequences can become embedded within modern environmental-response networks^26–31^.

Whether the same principle applies to hypoxia signalling and TE regulation, nevertheless, remains entirely unexplored. Most studies defining HIF1A targets rely on gene annotations or analyses that effectively exclude repetitive sequences. Consequently, we do not know if HIF1A actively recruits TE-derived regulatory sequences to coordinate immune and metabolic gene programs, which lies entirely outside the scope of traditional gene-centric analyses. Here, we systematically examine the relationship between HIF1A, hypoxia and the repetitive genome by analyzing genes and TEs within the same regulatory framework. We first used single-cell transcriptomic profiling of peripheral blood mononuclear cells (PBMC) exposed to hypoxia, pharmacological HIF stabilization and interferon stimulation to resolve how the gene and retrotranscriptome responses differ across immune populations. We then integrate HIF1A ChIP-seq, paired with transcriptomic data from multiple human cell systems to identify directly occupied genomic targets and determine whether particular TE families are preferentially recruited by HIF1A. We test whether these observations extend to primary immune cells from peripheral blood and tonsil, representing contrasting physiological oxygen environments, and use CRISPR deletion of individual TE loci to determine their consequences for host gene expression.

Across these complementary systems, we identify a striking asymmetry between the conventional transcriptome and the retrotranscriptome. Gene responses to hypoxia are highly cell-type dependent and distributed between activation and repression, whereas TE responses are overwhelmingly activating. Within this broader response, HIF1A engages a specific and reproducible subset of TE loci centred on LTR7/HERVH carrying canonical HREs, and loss of HIF1A DNA binding selectively affects the HERVH loci it occupies. Individual LTR7/HERVH loci can, in turn, modulate the expression of nearby genes *in cis* and distant targets *in trans*. These findings extend the canonical HIF transcriptional network into the repetitive genome and identify endogenous retroelements as a previously underappreciated component of the regulatory architecture through which human cells respond to oxygen.

## RESULTS

### Single-cell profiling reveals cell-type-specific gene response but a predominantly activating TE response to hypoxia

Circulating immune cells continually move between environments with markedly different oxygen availability, well-oxygenated blood to substantially more oxygen-restricted tissues and inflammatory sites^5^. We therefore began by asking how hypoxia reshapes gene and TE expressions across human immune-cell populations at single-cell resolution. PBMCs are therefore a physiologically appropriate starting point for our investigation. PBMCs from three healthy donors were cultured for 24 h under four conditions: normoxia (NOX), hypoxia (HOX, 1% O₂), pharmacological HIF stabilization with the prolyl hydroxylase inhibitor roxadustat (ROX), and interferon stimulation (IFN). Roxadustat stabilizes HIF1A without lowering oxygen, allowing us to distinguish responses associated with HIF activation from those specifically requiring low oxygen, whereas interferon provided an independent immune-stimulation control^32,33^. Cells were profiled by single-cell transcriptome and AbSeq protein sequencing on the BD Rhapsody platform, using a combined gene-and-TE reference^34^. After demultiplexing and quality control, cells were resolved into nine populations based on AbSeq and canonical RNA markers: CD4⁺ T cells, NK cells, monocytes, B cells, MAIT cells, neutrophils, basophils, plasmacytoid dendritic cells (pDCs), and a proliferating population (Figure 1B, C; Supplementary Figure S1).

Upon comparing the hypoxia with normoxia, we first asked whether all immune-cell populations responded similarly to reduced oxygen. Our analysis showed a unequal degree of transcriptome divergence across the nine cell types (Figure 1D and Supplementary Table 1). Monocytes were by far the most responsive population, with 3,010 significant differentially expressed genes (DEGs) under HOX versus NOX. They also showed the largest responses to roxadustat and interferon: 2,243 under ROX versus NOX; 936 under ROX versus HOX; and 3,752 under IFN versus NOX (p_adj_ < 0.05). In contrast, the hypoxic response was much smaller in CD4⁺ T cells (178 genes), NK cells (36), neutrophils (33), B cells (30), and MAIT cells (7), while basophils, pDCs, and proliferating cells showed little or no detectible response to either hypoxia or roxadustat (0 to 11 genes under HOX versus NOX).

We therefore asked whether these populations are also non-sensitive to IFN stimulation, as we found in the case of hypoxia. To address this, we compared the same cell populations following interferon stimulation, which are treated as an independent immune stimulus. Basophils, pDCs, and proliferating cells all mounted a detectable interferon response, with 16–51 significant DEGs under IFN versus NOX (Figure 1D and Supplementary Table S1), suggesting that these populations do respond to interferon under the same experimental and analytical set-up. Thus, their limited response is unlikely to be due to solely low cell numbers, although smaller hypoxia-induced changes may still fall below the detection threshold, as the statistical power depends on both size and cell abundances^35,36^.

The large differences in the number of DEGs raised the question of whether some immune populations fail to activate the canonical hypoxic response, or instead exhibit shallow transcriptional changes. We therefore examined established Hypoxia-inducible factors, e.g., HIF1A target genes across individual cell types. The expression of canonical HIF1A targets were induced by both hypoxia and roxadustat in the majority of lineages (Figure 1F). Fifteen genes were significantly induced (padj < 0.05) under HOX in at least three of the six cell types: monocytes, CD4⁺ T cells, NK cells, B cells, MAIT cells, and neutrophils. Key HIF-targets ENO1, GAPDH, GPI, LDHA, and P4HA1 in all six, whereas ALDOC, BNIP3L, PGK1, PKM, and TPI1 responded in five. Neutrophils showed the largest effects with a median log2 fold change of 2.81 across significant genes against 1.06 to 1.73 elsewhere. Among HIF-target genes, the largest change was observed for VEGFA at 3.02 in B cells (p_adj_ = 1.3 × 10⁻⁶), 2.99 in NK cells (5.2 × 10⁻¹⁴), and 2.85 in CD4⁺ T cells (1.1 × 10⁻⁵⁹). Interestingly, all fifteen genes behaved similarly under roxadustat (Figure 1F; Supplementary Figure S2A, Supplementary Table S1). Thus, despite the markedly different size of their overall transcriptional responses, the major immune-cell populations share a conserved and core HIF programme.

This shared HIF response was accompanied by the expected metabolic shift. Glycolysis increased across all eight annotated populations under both hypoxia and roxadustat, whereas mitochondrial respiration was suppressed only under hypoxia, indicating that this arm of the response requires reduced oxygen rather than HIF1A stabilization alone (Figure 1G; Supplementary Figure S2). Interferon stimulation, in turn, mounted the canonical interferon-stimulated gene response (Figure 1I and Supplementary Table S1). Roxadustat reproduced a substantial part of the hypoxic transcriptional programme, though the overlap varied by population (Jaccard similarity 0.26-0.51; Supplementary Figure S3), with monocytes showing the largest hypoxia-specific component (936 DEGs; Figure 1H; Supplementary Table S1).

### Transposable elements reveal a strikingly different response

We sought to understand if the TEs followed the similar pattern of expression as genes. Strikingly, in contrast to gene expression changes, the retrotranscriptome behaved in a different but in a consistent way. While gene responses varied extensively between cell types and were distributed between activation and repression, differentially expressed TE families were overwhelmingly activated across all three perturbations (Figure 1D-E and Supplementary Figure S3A-F). Of 140 TE families responding significantly to hypoxia, 135 (96%) increased in expression. Similar responses were noted under roxadustat, with 84 of 90 (93%), and interferon stimulation, with 53 of 61 (87%) were induced. In contrast, only 51%, 41%, and 51% of differentially expressed genes were upregulated under hypoxia, roxadustat, and interferon, respectively.

This pronounced activation bias was not an inherent feature of TE quantification or differential-expression testing. When the two stimulated conditions, roxadustat and hypoxia were compared directly, TEs changes became bidirectional: 38% of significant TE families were increased under roxadustat, matching closely with 40% of observed genes (Figure 1E). Together, despite the different pattern of gene expression programs following the distinct stimulations viz. hypoxia, HIF stabilization and interferon, the TE families responding to these stimulations are overwhelmingly unidirectional as they are activated rather than repressed.

Together, the single-cell analysis revealed two fundamentally different features of the hypoxic response. Gene regulation was highly dependent on immune-cell identity and included both activation and repression, whereas the TE compartment showed a striking and reproducible bias toward increased expression. The same directional pattern was observed after pharmacological HIF stabilization and, to a lesser extent, interferon stimulation, suggesting that TE activation represents a broader feature of cellular stress responses. However, the limited sequencing depth of single-cell profiling restricted our ability to resolve individual responsive TE loci.

### Hypoxia drives an activating TE response in primary immune cells from blood and tonsil

Having identified a predominantly activating TE response to hypoxia across PBMC populations, we next asked whether this behaviour could be reproduced by deeper RNA sequencing of primary immune cells and whether it varied according to their physiological context. We compared cells derived from peripheral blood and tonsil, representing markedly different oxygen environments, because circulating blood is relatively well oxygenated whereas lymphoid tissues, including tonsillar germinal centres, exist under substantially lower oxygen enviroment. Primary immune cells from blood (three donors) and tonsil (four donors) were cultured under normoxia or hypoxia, either at rest or following TCR stimulation with anti-CD3/CD28 beads, thereby allowing us to examine the interaction between oxygen availability, tissue origin, and activation state (Figure 2A; Supplementary Table S2).

**Figure 2.**
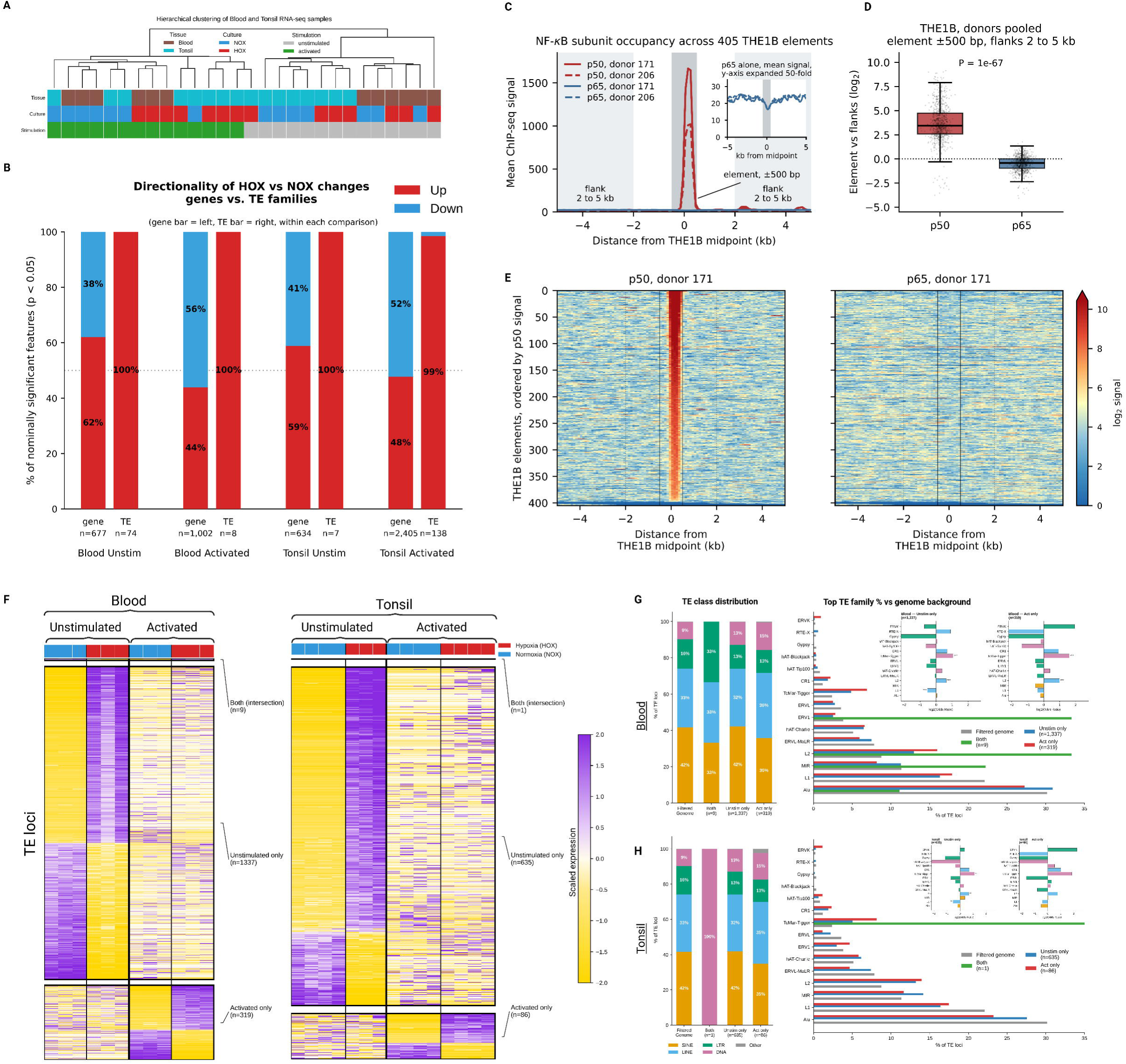
Hypoxia activates transposable elements in primary immune cells from blood and tonsil. **(A)** Hierarchical clustering of all blood and tonsil-derived immune cell RNA-seq samples based on gene and TE expression. Annotation bars indicate tissue, hypoxia/normoxia condition, and stimulation state. **(B)** Directionality of hypoxia versus normoxia changes for genes and TE families in each of the four tissues and activation contexts. Within each comparison, the gene bar is on the left and the TE bar on the right. “n” denotes the number of features. TE responses are induced to 99%-100%. **(C)** Mean NF-κB ChIP-seq signal for the p50 and p65 subunits across a 10 kb window centred on 405 THE1B elements, which have peaks on those loci, in activated human CD4⁺ T cells from two donors. The shaded band marks the element footprint. The inset replots the p65 tracks on a 50-fold expanded y-axis. **(D)** Enrichment of each element over its own flanking regions. The element is defined as ±500 bp from the midpoint and the flank is defined as 2 to 5 kb on both sides, donors are pooled. P-value is calculated by the paired Wilcoxon signed-rank test. **(E)** Per-element ChIP-seq signal for p50 (left) and p65 (right) in donor 171, elements ordered by p50 signal in the element and the same order used for both panels. The colour scale is shared between panels. **(F)** Locus-level TE expression in blood (left) and tonsil (right), split by stimulation context. Rows are individual TE loci grouped into those passing in both stimulation states, in unstimulated cells only, and in activated cells only. **(G)** TE class and family composition of the responsive loci in blood relative to the mapping background. On the left, class distribution is shown. On the right, the most frequent families compared with the genome background are shown, and the inset shows log₂ odds ratios. Filtered genome corresponds to the filtered TE annotations for the mapping, see the method for genome index preparation. **(H)** The same analysis for tonsil, in the same way as in (G).

We first confirmed that primary blood- and tonsil-derived immune cells mounted a canonical hypoxic response. The core HIF1A glycolytic program was induced in all four comparisons, with the strongest response in unstimulated tonsil-derived cells and a more modest but concordant response in blood-derived cells (Supplementary Figure S4; Supplementary Table S2). Canonical targets including MIR210HG, VEGFA, PFKFB4, ALDOC, GPI, ENO1, and PGK1 were consistently induced. TCR activation, however, changed the metabolic response to hypoxia in a selective manner. In activated blood-derived cells, SLC2A1/GLUT1, SLC2A3, and HK2 were reduced under hypoxia, whereas none showed this behaviour in resting cells, validating the reliance of our experimental set-up (Supplementary Figure S4, S5, and Supplementary Table S2). Besides, the key immune effector genes were upregulated under hypoxia (Supplementary Figure S6A,B).

We then asked whether the activation bias of TEs observed in single-cell PBMCs was preserved across tissue origin and activation state. Among nominally significant TE families, every responsive family was induced in unstimulated blood, activated blood, and unstimulated tonsil, and 136 of 138 were induced in activated tonsil (Figure 2B). The same directional bias remained when all tested families were considered irrespective of significance, with 69–93% shifting upward across the four comparisons. Genes showed no similar directional behaviour, remaining approximately balanced between induction and repression in the same samples. Activated tonsil provided the strongest signal: after FDR correction, 53 TE families remained significant, and all 53 were induced (Figure 2B and Supplementary Table S2). Thus, the predominantly activating TE response is not restricted to PBMC single-cell data. It is reproduced by deeper bulk sequencing in primary human immune cells and persists across two tissue environments and distinct activation states. Of note, the number of significantly differentially expressed genes, not only TEs, scaled with activation state within each tissue (blood: 38 unstimulated versus 216 activated; tonsil: 70 versus 885; padj < 0.05), and the direction of the response shifted with it, from predominantly induction in resting cells to predominantly repression after the activation (Supplementary Table S2 and Supplementary Figure S6C).

The strong family-level activation bias tempted us to ask if the same TE loci responsive in resting and activated cells, or does cellular state determine which copies are upregulated? Locus-level analysis revealed remarkably little overlap. Only 9 of 1,665 responsive loci in blood and 1 of 722 in tonsil were shared between resting and activated conditions (Figure 2F). Moreover, the responsive loci were drawn non-randomly from the repeat landscape and differed in class and family composition between resting and activated cells (Figure 2G,H). Thus, hypoxia consistently pushes the retrotranscriptome toward activation, but the identity of the elements responding is strongly determined by cellular state.

THE1B provided an example of this context-dependent TE regulation. THE1B showed an upward trend under hypoxia in all four primary-cell comparisons and reached the highest level in activated tonsil cells (Supplementary Figure S5 and Supplementary Table S2). Because THE1B elements contain NF-κB-responsive sequences^37,38^, we examined published NF-κB ChIP-seq from activated human CD4⁺ T cells and observed strong p50 occupancy centred on these elements (Figure 2C–E, Supplementary Figure S5). The selective induction of THE1B in activated tonsil therefore suggests that hypoxia-responsive TE regulation can intersect with activation-dependent immune transcription-factor networks.

Together, these data show that hypoxia produces a remarkably consistent directional effect on the primary immune-cell retrotranscriptome while the identity of the responsive elements is strongly shaped by tissue and activation state.

### Hypoxia induces balanced gene regulation but directional transposable-element activation across human cell lines

Our results from single-cell PBMCs and in primary blood- and tonsil-derived immune cells suggested that unidirectional TE activation may represent the cellular response to low oxygen. To test this possibility in an independent experimental context, and to obtain responsive TE families more robustly, we analyzed bulk RNA-seq from four human cancer cell lines A549, H460, HeLa, and RKO, cultured under matched normoxic and hypoxic conditions. Gene and TE expression were quantified simultaneously against the same combined reference used throughout the study. Unsupervised hierarchical clustering separated hypoxic from normoxic samples in every cell line, confirming a coherent transcriptional response to low oxygen (Figure 3A).

**Figure 3.**
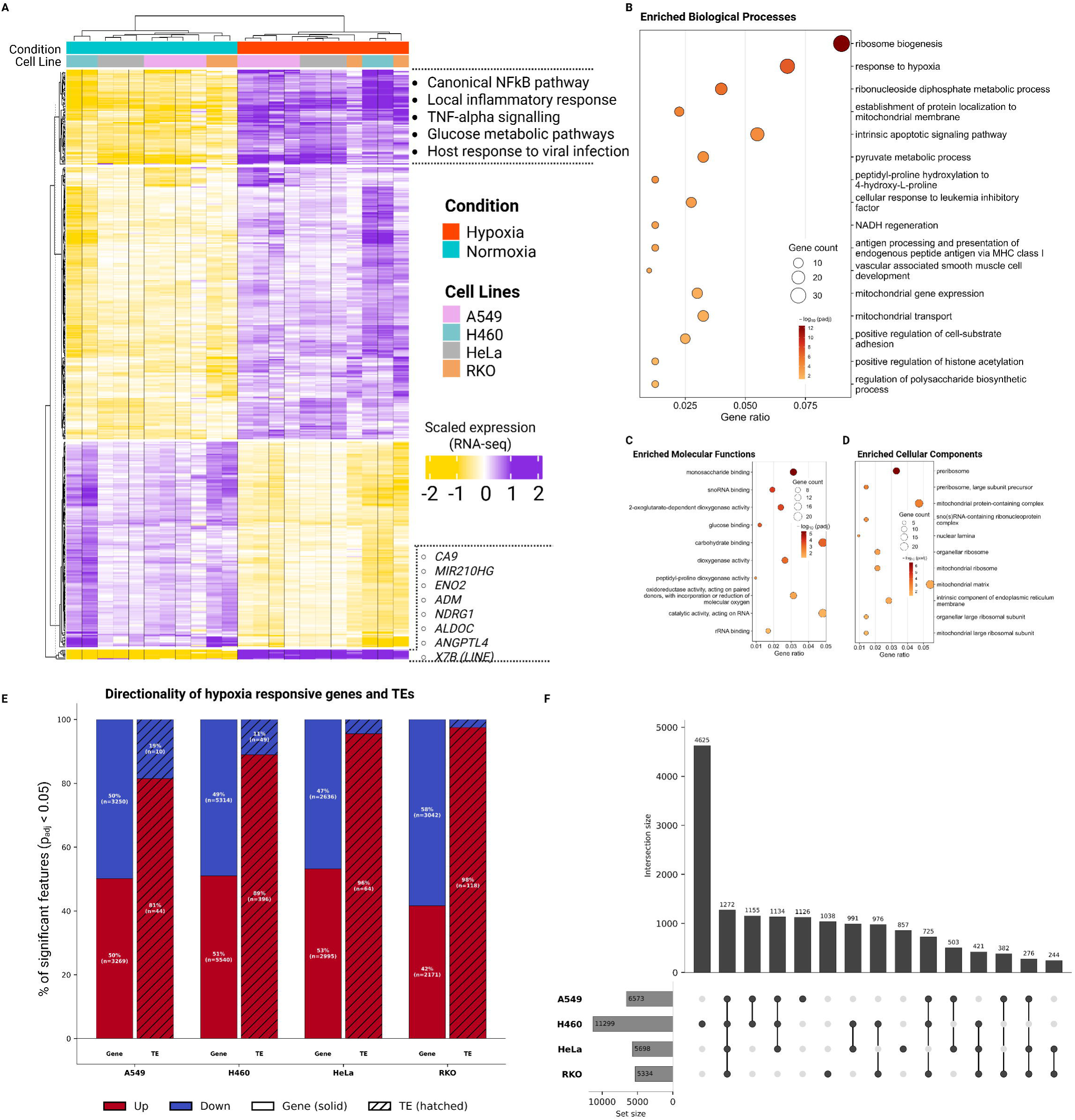
Hypoxia produces a bidirectional gene response and a near-unidirectional TE response in human cancer cell lines. **(A)** Unsupervised hierarchical clustering and heatmap of scaled RNA-seq expression for significantly differentially expressed genes and TEs across four transformed (cancer cell) lines (A549, H460, HeLa, RKO) under normoxia and hypoxia. Annotation bars indicate condition and cell line. Pathways over-represented in the hypoxia-induced block are listed at the right, and some of the hypoxia-responsive features are indicated at the lower right, including the canonical targets CA9, MIR210HG, ENO2, ADM, NDRG1, ALDOC and ANGPTL4 together with the LINE family X7B. **(B)** Gene ontology over-representation analysis of differentially expressed genes for biological process terms. Bubble size reflects the number of annotated genes and colour reflects the adjusted p-value. **(C)** Enriched Molecular Function terms, plotted in the same way as in (B). **(D)** Enriched Cellular Component terms, plotted in the same way as in (B). **(E)** Directionality of the hypoxic response in each cell line, shown as the percentage of significant features induced (red) and reduced (blue) for genes (solid) and TE families (hatched). **(F)** UpSet plot of differentially expressed genes shared between cell lines. Bars show intersection sizes, and the left panel shows the total set size per line.

We first confirmed that each line overexpressed a canonical hypoxic transcriptional response. Canonical HIF1A targets, including CA9, ALDOC, MIR210HG, ENO2, and ADM were strongly induced across all four lines, and 385 genes were significantly upregulated in every cell line (Figure 3A, Supplementary Figure S7A, Supplementary Table S3), although many DEGs were line-specific (Figure 3F). These shared genes were enriched for the expected hypoxia-associated programmes, including glycolysis and pyruvate metabolism^39^, but also for inflammatory and stress-response pathways involving NF-κB, TNF signalling, and host responses to viral infection (Figure 3A–D). Thus, despite their different cellular origins, the four lines shared a conserved core hypoxic programme but also illustrated cell-line-specific transcriptional responses.

Across all four cell lines, DEGs remained approximately balanced between induction and repression. In contrast, 81–98% of significantly responsive TE families were induced under hypoxia (Figure 3E; Supplementary Table S3). In RKO cells, for example, 118 TE families’ expression was increased, and only three were reduced (Figure 3E and Supplementary Table S3). The unidirectional TE response, therefore, is reproduced across diverse transformed human cell types. A clear pattern emerged among LTR-class elements, including LTR7, LTR7B, LTR7C, LTR7Y, and HERVH-int, which ranked top among the most significant TE families in every analyzed line (Supplementary Figure S7B). The effect was particularly pronounced in H460, where both LTR7 and HERVH-int were significantly upregulated. The recurrence of LTR7/HERVH across independent cell lines pointed to it as a key candidate of the hypoxic TE response.

Together, these data extend the near-unidirectional TE response observed from primary immune cells to unrelated cell lines and identify LTR7/HERVH as a key hypoxia-responsive family. Expression alone, however, cannot distinguish whether these elements are direct HIF1A targets or are activated indirectly as part of the broader hypoxic programme. We therefore turned to genome-wide HIF1A occupancy to understand if its direct binding precedes the transcriptional outcomes of HERVH.

### Genome-wide HIF1A occupancy recovers the canonical hypoxic regulatory network

The transcriptional analyses identified a broad hypoxic response and repeatedly implicated LTR-class elements. We therefore assembled 17,019 HIF 1A ChIP-seq peaks from A549, H460, HCT116, RKO cells profiled under hypoxia and normoxia, and mapped them simultaneously to gene and TE annotations (Figure 4A, B and Supplementary Table S4). Before interrogating the repetitive genome, we first asked whether this catalogue recovered the established genomic architecture of HIF1A regulation. 44.6% of peaks fell at promoters, 29.3% within gene bodies and 15.8% in intergenic regions, while 10.3% overlapped TE-derived sequence (Figure 4B and Supplementary Table S4).

**Figure 4.**
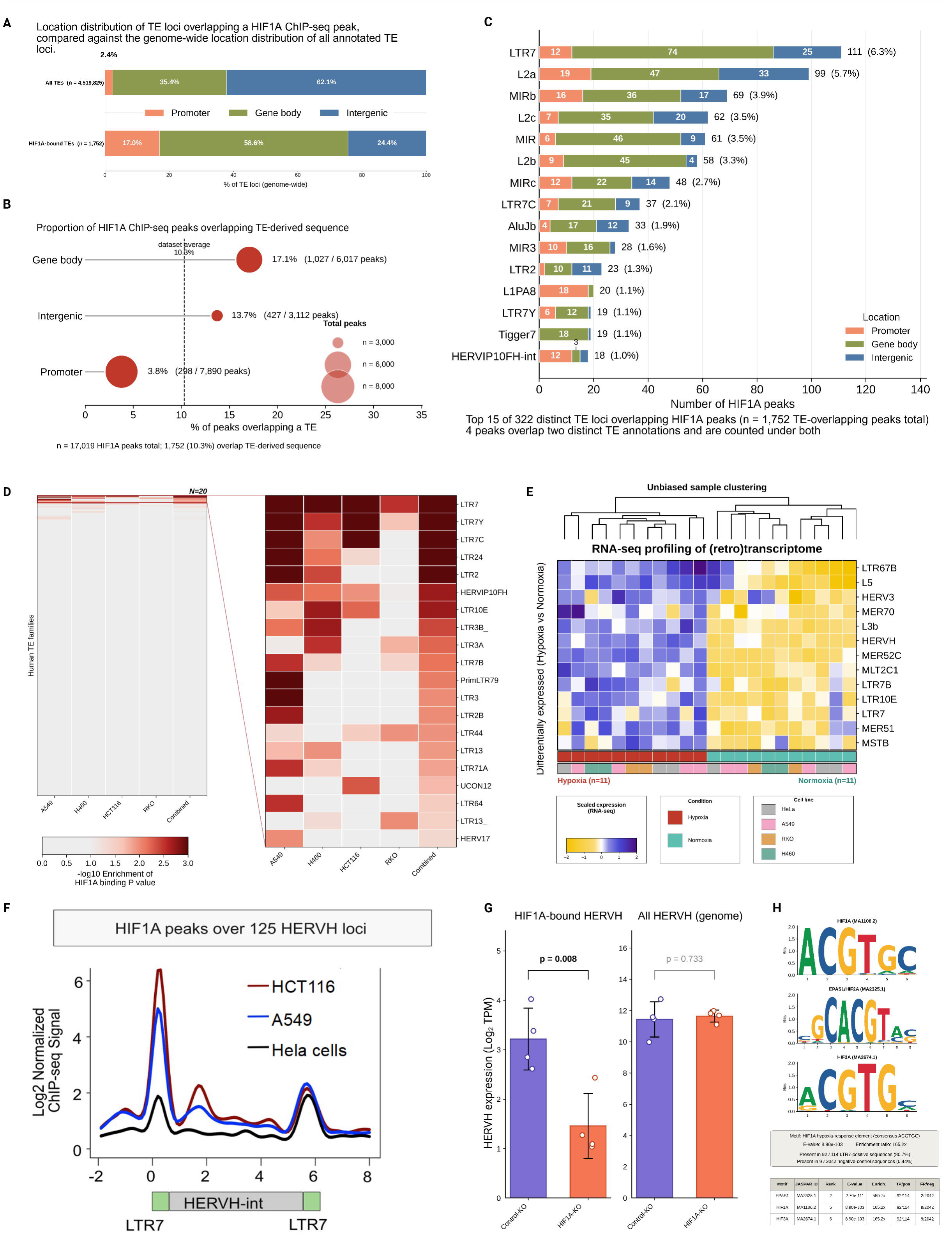
The same HIF1A target catalogue, applied to transposable elements, converges on LTR7/HERVH. **(A)** Genomic context of TE loci overlapping a HIF1A ChIP-seq peak compared with the genome-wide distribution of all annotated TE loci. **(B)** Proportion of HIF1A peaks overlapping TE-derived sequence in each genomic context. Bubble size reflects the number of peaks per category, and the dashed line shows the dataset average, which is 10.3%. **(C)** The 15 most frequently HIF1A-bound TE families, stacked by genomic location of the overlapping peak. Binding falls into 322 distinct families across 1,752 TE-overlapping peaks; LTR7 ranks first with 111 peaks. **(D)** Permutation-based enrichment of HIF1A binding at each TE family (regioneR, 1,000 permutations). The enrichment is tested separately in each cell line and all together. The expanded panel shows the 20 most enriched families. LTR7 and LTR7 subfamilies like LTR7Y are the top-enriched families by HIF1A binding. **(E)** Heatmap of scaled RNA-seq expression for differentially expressed TE families across hypoxic and normoxic samples, with unbiased sample clustering above and condition and cell line annotation below. **(F)** Mean HIF1A ChIP-seq signal across 125 full-length HERVH loci in the genome using HCT116, A549 and HeLa cells. The x-axis spans the provirus with the LTR7-HERVH-int-LTR7 structure indicated below. Signal peaks sharply over the flanking LTR7 regions. **(G)** Aggregated HERVH expression in HIF1A knockout versus control cells. Left: HERVH loci overlapping a HIF1A ChIP-seq peak are significantly reduced in the knockout (p = 0.008). In the right bar plot, all genome-wide HERVH loci are unchanged (p = 0.733). Points are biological replicates, and error bars show standard deviation. **(H)** Sequence motifs enriched within HIF1A-bound LTR7 sequences relative to unbound LTR7 loci, identified by MEME-SEA against JASPAR 2026. The canonical HIF1A motif (MA1106.2, consensus ACGTGC) is present in 80.7% of bound copies, compared with 0.44% of unbound copies, a 165-fold enrichment.

We then tested whether our analysis recovered known HIF1A-responsive pathways. HIF1A signal increased prominently at the promoters of immune-response and immediate-early genes (IEGs) under hypoxia compared with normoxia (Figure 5A; Supplementary Figures S9 and S10). Among the most reproducibly occupied immune loci were VEGFA, RELA, NFIL3, and ADM, while IEG targets included JUN, FOSB, FOSL2, DDIT4, and DUSP5. Thus, our catalogue captures both the canonical metabolic response to low oxygen and its intersection with stress- and immune-responsive genesets.

**Figure 5.**
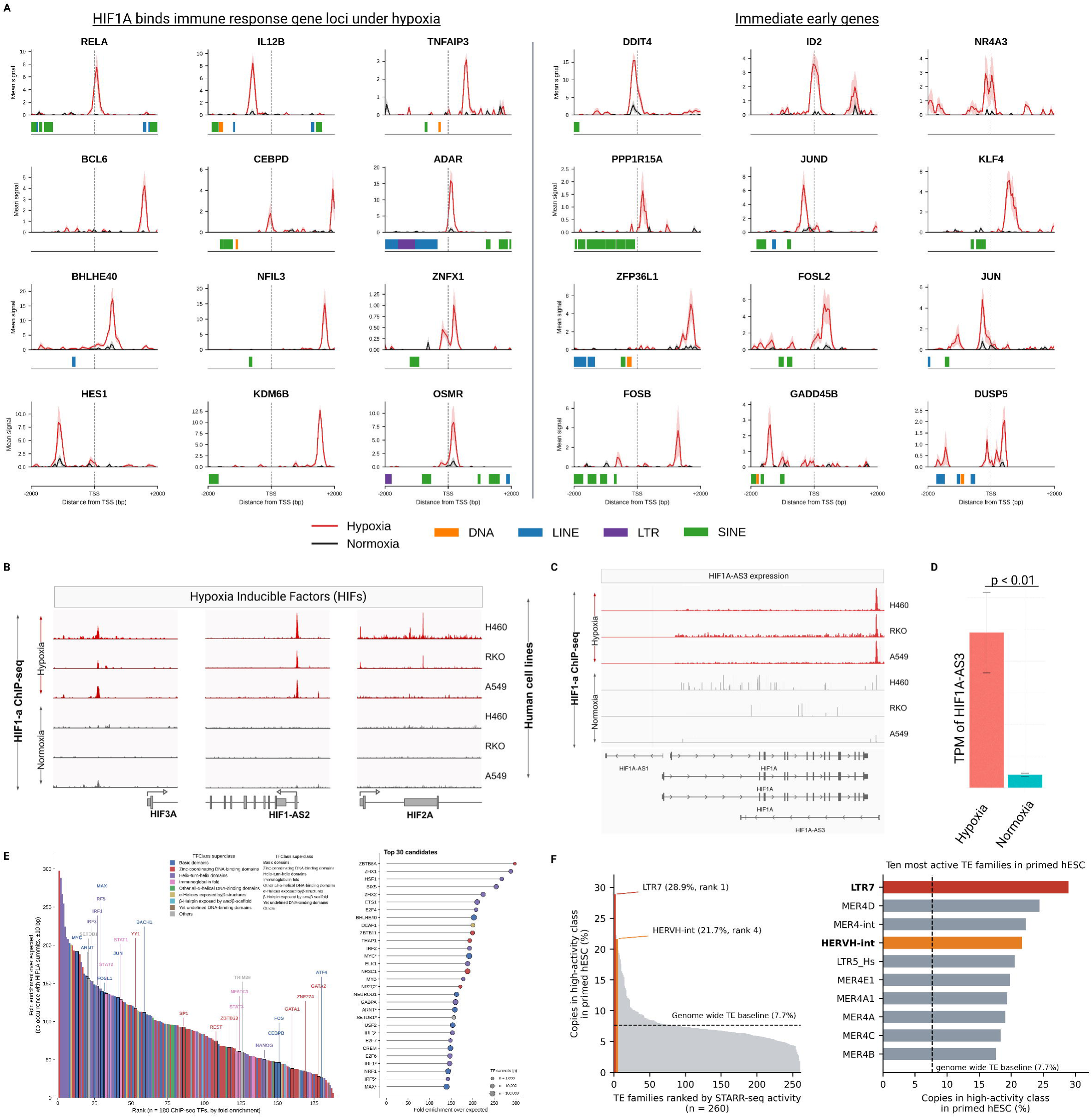
Genome-wide mapping of HIF1A occupancy recovers the established gene targets. **(A)** HIF1A ChIP-seq signal in a ±2 kb window around the transcription start site for representative immune response genes (left) and immediate early genes (right), under hypoxia (red) and normoxia (black). TE loci within each window are annotated by class below the x-axis. **(B)** Genome browser tracks of HIF1A ChIP-seq signal at the HIF3A, HIF1A-AS2, and HIF2A loci under hypoxia and normoxia. **(C)** Genome browser tracks at the HIF1A-AS3 locus. A strong hypoxia-specific peak is present in all cell lines. Gene models for HIF1A-AS1, HIF1A, and HIF1A-AS3 are shown below. **(D)** HIF1A-AS3 expression (TPM) under hypoxia versus normoxia, pooled across cell lines (p < 0.01, two-sample Student’s t-test). Error bars show standard deviation. **(E)** Co-localisation of HIF1A summits with 188 curated ENCODE transcription factor ChIP-seq datasets. On the left, all factors ranked by fold enrichment over expected co-occurrence, coloured by TFClass superclass. On the right, the 30 most enriched factors, with dot size reflecting the number of summits per factor. **(F)** ChIP-STARR-seq activity of TE families in primed human embryonic stem cells. On the left, all 260 tested families ranked by the percentage of copies in the high-activity class, and the genome-wide TE baseline is 7.7%, which is indicated by a dashed line. LTR7 ranks first at 28.9% and HERVH-int fourth at 21.7%. On the right, the ten most active families are shown.

Leveraging our catalogue, we asked if HIF1A also regulates components of its own network. We found that HIF1A also occupied the components of the HIF family members itself, suggesting that the response contains regulatory feedback. Hypoxia-specific peaks were detected at HIF1A-AS2, HIF1A-AS3, HIF2A, and HIF3A in multiple lines (Figure 5B, C). HIF1A-AS3 showed reproducible binding across lines, and was correspondingly induced under hypoxia (Figure 5C-D). Because antisense transcription at the HIF1A locus can suppress HIF1A *in cis* by impeding transcriptional elongation and H3K4me3 deposition^40^, direct HIF1A recruitment to HIF1A-AS3 places HIF1A within a potential negative feedback circuit controlling its own activity.

We then asked which transcription-factor networks align with HIF1A-bound regulatory regions. HIF1A summits were compared with 188 curated ENCODE TF ChIP-seq datasets, and because all factors showed significant co-localization after multiple-testing correction, we used enrichment magnitude to identify the most informative associations (Figure 5E). The strongest interpretable associations included the HIF partner ARNT and other bHLH/bZIP factors such as BHLHE40, MYC, JUN, FOS and CEBPB. HIF1A sites were also strongly enriched for IRF and STAT occupancy, including IRF1/2/3 and STAT1/2/3, linking HIF1A-bound regions to the immune and inflammatory programmes identified in the RNA-seq data. These results position HIF1A within a broader stress-responsive regulatory network in which oxygen sensing intersects with AP-1, interferon and cytokine signalling (Supplementary Table S7).

Together, these analyses establish that the HIF1A occupancy catalogue recovers the expected architecture of oxygen-responsive gene regulation, including canonical HIF targets, feedback within the HIF pathway and alignment with immune and stress-responsive transcription factors.

### HIF1A selectively targets LTR7/HERVH endogenous retroviral elements

Having established that the HIF1A targets its expected gene targets in our analysis, we next asked whether HIF1A binds TEs sequences randomly or is enriched in particular TEs. Of the 17,019 HIF1A peaks, 1,752 (10.3%) overlapped TE-derived sequence (Figure 4A,B). These TEs-associated peaks were not distributed uniformly.17.1% of peaks are in gene bodies (1,027 of 6,017), and 13.7% of peaks are intergenic (427 of 3,112), and 3.8% peaks are promoter peaks (298 of 7,890) overlapping with a TE (Figure 4B). HIF1A-bound TEs were strongly shifted toward gene-proximal positions, with promoter associated elements rising from 2.4% of the genomic TE background at promoters to 17.0% of HIF1A-bound TEs, while intergenic elements decreased from 62.1% to 24.4% (Figure 4A). This suggest that HIF1A preferentially engages a gene-proximal subset of the repetitive genome rather than sampling TEs according to their genomic abundance.

We therefore asked whether HIF1A occupancy was specific to particular TE families. Among 322 distinct TE families with HIF1A peaks, the single most frequently bound family was LTR7 with 111 overlapping peaks (Figure 4C). Related LTR7C and LTR7Y were also among the most frequently bound families. Together, these annotations immediately pointed to the LTR7/HERVH endogenous retroviral group.

To distinguish preferential HIF1A recruitment from TE abundance, we tested the enrichment for each family^41^ against a matched random background by permutation. LTR7 was significantly enriched in every individual cell-line and in the combined HIF1A peak set, while the related LTR7C and LTR7Y subfamilies ranked too within the same enriched repertoire of TEs (Figure 4D; Supplementary Table S5). LTR7 is the long terminal repeat of HERVH, with the 51 LTR functioning as the promoter of the provirus; LTR7C and LTR7Y represent related variants of this regulatory sequence^42^. Overall, two independent observations point to the one TE family: LTR7/HERVH enrichment in HIF1A bindome, and was induced by hypoxia at the RNA level (Figure 4E; Supplementary Table S3).

We next asked where HIF1A binds within the HERVH provirus. Averaging HIF1A ChIP-seq signal across 125 full-length HERVH loci revealed a sharp maximum over the flanking LTR7 elements and substantially weaker signal across the internal HERVH sequence (Figure 4F). Because the 51 LTR7 constitutes the HERVH promoter, HIF1A occupancy is localised at the regulatory element that drives HERVH transcription.

We therefore examined hypoxic HERVH expression in HIF1A-deficient HCT116 cells. To establish whether HIF1A is required for HERVH expression, we examined HCT116 HIF1A−/− cells, in which exons 3 and 4 of the HIF1A were deleted jeopardizing all three reading frames^43^. HERVH loci overlapping HIF1A ChIP-seq peaks showed significantly and dramatically lower expression in HIF1A-KO cells (p = 0.008, two-sample Student’s t-test). In contrast, expression summed expression across all 1,695 detectable HERVH loci, was unchanged (p = 0.733; Figure 4G). The effect is thus restricted to HIF1A occupied loci, providing a functional support for the direct and locus-specific activation.

The enrichment of LTR7 compelled us to ask what distinguishes bound from unbound elements. We therefore compared sequences surrounding HIF1A-bound LTR7 loci with unbound LTR7 copies for transcription-factor (TF) motifs. We performed motif enrichment using MEME-SEA^44^ on ±50 bp windows around HIF1A summits. We found that the canonical HIF1A/HRE motif was present in 80.7% of bound LTR7 sequences but only in 0.44% of unbound sequences, representing approximately 165-fold enrichment (Figure 4H; Supplementary Table S6). Related HIF/ARNT-family motifs built around the same HRE core were similarly enriched. HIF1A binding to LTR7 is therefore strongly sequence-directed: among thousands of related LTR7 copies, HIF1A preferentially occupies those carrying an embedded HRE.

Sequence recognition explains where HIF1A binds, but not whether the bound elements are competent regulatory sequences. Ranking 260 TE families by ChIP-STARR-seq^45^ activity in primed human embryonic stem cells placed LTR7 first, with 28.9% of copies in the high-activity class, and HERVH-int fourth at 21.7%, against a genome-wide TE baseline of 7.7% (Figure 5F). LTR7/HERVH is therefore both the preferred HIF1A substrate among repeats and one of the most autonomously active regulatory families in the repetitive genome.

We finally asked whether the loss of HIF1A binding abolishes TE response globally in response to hypoxia. Even in HIF1A-deficient cells, hypoxia induced 349 TE loci while only one was reduced (padj < 0.05) (Supplementary Figure S8). In sum, our analysis supports that HIF1A is required for the direct activation of LTR7/HERVH loci it binds, but it is not required for the TE activation as a whole. Loss of HIF1A instead redirects the hypoxic retrotranscriptome toward an alternative repertoire, revealing HIF1A-dependent routes of TE activation.

### CRISPR deletion establishes regulatory activity of HIF1A-bound LTR7/HERVH loci

HIF1A binding to HRE-containing LTR7/HERVH elements suggested that these loci may be acting as cis-regulatory elements and contribute to re-wiring neighbor gene expression and to overall physiology. We therefore tested two individual HIF1A-bound LTR7/HERVH elements for their potential to contribute host gene regulation. Two full-length loci overlapping HIF1A ChIP-seq peaks were selected for CRISPR deletion and are hereafter referred to as TG1 (*chr19:15,828,728-15,829,215; hg38*) and TG2 *chr5:147,869,835-147,870,673; hg38*) (Figure 6; for HIF1A ChIP-seq peaks: Supplementary Table S4). Because the HERVH/LTR7 activity is known to maintain human embryonic stem cells and these two loci are actively expressed, we chose human embryonic stem cells to perform our CRISPR study. RNA-seq of knockout and control cells was then used to determine the transcriptional consequences of removing each element.

**Figure 6.**
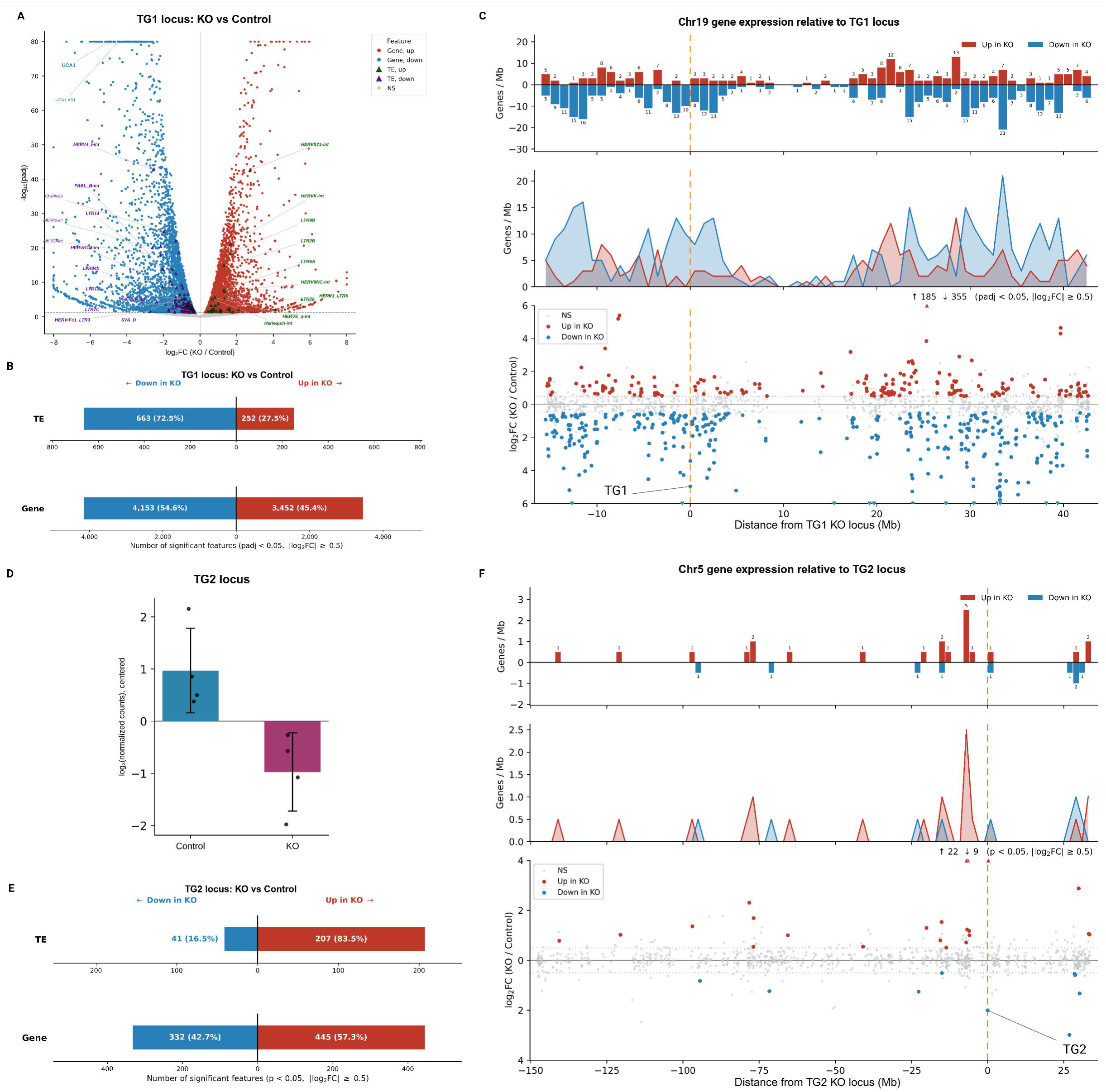
CRISPR deletion demonstrates locus-specific regulatory footprints at LTR7/HERVH elements. **(A)** Volcano plot of differentially expressed features following deletion of the TG1 locus, genes and TE loci indicated separately. **(B)** Bar plot showing the number of differentially expressed TE loci and genes, grouped by direction of change (up- or downregulated), within the TG1 locus in KO versus control. (padj < 0.05, |log₂FC| ≥ 0.5). **(C)** Chromosome 19 gene and TE expression relative to the TG1 locus. On top, the density of significantly changed genes per Mb is shown. In the middle, line profiles of the significantly changed genes are shown. On the bottom, log₂ fold change for all genes with distance information from the deleted locus is shown. The closest gene to the deleted TG1 locus is labelled as TG1. **(D)** Centred log₂ normalised counts for the closest gene to the TG2 locus in control and knockout cells. Points are replicates, and error bars show standard deviation. **(E)** Bar plot showing the number of differentially expressed TE loci and genes, grouped by direction of change (up- or downregulated), within the TG2 locus in KO versus control. (p < 0.05, |log₂FC| ≥ 0.5). **(F)** Chromosome 5 gene and TE expression relative to the TG2 locus. On top, the density of significantly changed genes per Mb is shown. In the middle, line profiles of the significantly changed genes are shown. On the bottom, log₂ fold change for all genes with distance information from the deleted locus is shown. The closest gene to the deleted TG2 locus is labelled as TG2.

Deletion of TG1 (Supplementary Figure S11A) produced a broad transcriptional response, altering the expression of thousands of genes and TEs (Figure 6A, B, Supplementary Figure S11B, Supplementary Table S8). Importantly, the effect was stronger in the genomic window around the deleted locus. Of 43 expressed genes located within 1 Mb of TG1, 22 were DEGs among which the 18 showed the reduced expression (Figure 6C and Supplementary Table S8). The predominance of decreased expression extended across a substantial region surrounding the deletion, indicating that TG1 contributes to the activity of a broader local regulatory domain rather than controlling only a single proximal transcript (Figure 6C).

TG2 (Supplementary Figure S11C) produced a smaller but distinct regulatory scenario. At FDR significance, deletion altered 27 genes together with 11 TE loci, while a broader response was evident at nominal significance (Figure 6D, E, Supplementary Figure S11D, Supplementary Table S9). The transcript immediately associated with the locus showed a marked reduction after TG2 deletion (Figure 6D, F), although variability between knockout replicates prevented this change from reaching FDR significance (Supplementary Table S9). In parallel, several HERVH loci elsewhere in the genome increased in expression, raising the possibility that removal of one active HERVH element can influence the broader retrotranscriptomic network.

Thus, two independently selected LTR7/HERVH loci identified solely by HIF1A occupancy produced measurable transcriptional output when deleted. The magnitude of these effects differed between TG1 and TG2, consistent with the regulatory output of individual retroelements being shaped by their genomic context. Together with the sequence-specific HIF1A occupancy and HIF1A-dependence established above, these CRISPR experiments provide functional evidence that HIF1A-bound LTR7/HERVH elements can act as regulatory components of the host genome, but they do not yet resolve the physiological consequences of this regulation or identify the specific neighbouring genes contacted directly by these elements. Future studies combining locus-specific perturbation with chromatin-conformation mapping and physiological readouts will be required to define their promoter interactions and determine how these regulatory circuits influence cellular adaptation to hypoxia.

## DISCUSSION

In this study, we asked whether HIF1A acts on TEs when oxygen availability is limited, and if so, which ones, through what mechanism, and with what consequence. Our findings extend the canonical mammalian response to oxygen beyond the gene-centric genome. Hypoxia produced a markedly different response in genes and TEs: while gene regulation remained bidirectional and strongly dependent on cellular identity, the retrotranscriptome was consistently biased toward activation. Within this broader response, HIF1A recognizes HRE motifs embedded within LTR7/HERVH copies that were carried into the primate germline by retroviral infection around 40 million years ago^46,47^, and has repurposed these ancient sequences as components of the canonical hypoxic transcriptional program. Many TEs still can respond to hypoxia without being bound by HIF1A. How strongly and in which direction a TE responds depends on the cell type and/or its condition, such as immune-activated cell states. And when we delete individual TE loci, what changes in the surrounding genome depend on where that locus is located, not on which TE family it belongs to. These findings identify endogenous retroviral sequences as components of the regulatory architecture through which human cells respond to oxygen availability.

### HIF1A is a sequence-directed activator of LTR7/HERVH elements

Roughly 80% of HIF1A-bound LTR7 copies carry the canonical HRE core sequence, while fewer than 1% of unbound copies do. It is site-specific recognition of an embedded motif. The HERVH loci overlapping HIF1A peaks are selectively reduced in expression when HIF1A loses DNA-binding capacity, while the remaining HERVH-int loci remain unaffected (Figure 4G). This shows that HIF1A directly switches on the HERVH loci it binds. The position of HIF1A binding within LTR7/HERVH also matters mechanistically. Because LTR7 serves as the promoter long terminal repeat for HERVH, the binding of HIF1A to the LTR7 locus positions the transcription factor directly at the regulatory element required to drive downstream HERVH transcription. The resulting transcripts, that is, HERVH transcripts, include long non-coding RNAs with documented trans-regulatory activity^48–51^, meaning that activation of a single LTR7 locus can influence gene expression beyond the immediate neighbourhood. This fits a broader principle emerging from ERV co-option: LTRs evolved to recruit cellular transcription factors and initiate transcription; after endogenization, the same properties make them unusually suitable substrates for regulatory innovation. HIF1A recruitment to LTR7/HERVH extends this principle to oxygen sensing.

The HIF1A-KO data support this: loss of HIF1A binding does not globally suppress TE transcription but shifts the upregulated TE repertoire toward a largely different, HIF1A-independent set of elements, suggesting that TE transcription under hypoxia is a net activating process driven by multiple parallel mechanisms, of which HIF1A-mediated activation of LTR7/HERVH is the most reproducible and sequence-specific under wild-type conditions. HIF1A is therefore exploiting HERVH at the same sequence class that has retained the HRE and originally evolved to control retroviral transcription. This provides a simple route for regulatory innovation: instead of evolving individual HIF-responsive elements independently, insertion and subsequent retention of retroviral LTRs could distribute pre-existing transcription-factor recognition sequences across the genome.

### TEs as active regulatory elements in response to hypoxia

A second feature of the response was its directionality. The unidirectional TE response to hypoxia is a distinct phenomenon from the gene response. In single-cell PBMC data, the TE responses to hypoxia and roxadustat were overwhelmingly induced. If TE activation under hypoxia were a non-specific consequence of global chromatin relaxation or transcriptional noise, we would expect a roughly equal mixture of induction and repression. Instead, across every system we examined, viz., single-cell PBMCs, primary blood and tonsil cells, and four cancer cell lines showed 68–98% of responsive TE families shifted upward. This reproducible activation skew is specific to TEs and contrasts sharply with the balanced bidirectional gene response measured in the same samples.

The most straightforward explanation is that HIF1A functions primarily as a transcriptional activator^52^ and because many TE copies carry strong internal promoters inherited from their retroviral ancestors^53,54^, the balance of chromatin effects at TE loci under hypoxia may be more consistently activating than at gene loci, where repressive mechanisms and competing transcription factors produce the bidirectional pattern we observe. One possibility is that many responsive retroelements retain autonomous or semi-autonomous regulatory sequences that convert stress-responsive transcription-factor activity preferentially into transcriptional gain.

Our findings provide an example of molecular exaptation in which retroviral regulatory sequences have been incorporated into a host environmental-response network. The transcription-factor binding sites and promoter activity that once served viral replication did not disappear on endogenization; they became raw material available to host regulatory networks^53,54^. In this case, HRE-containing LTR7 copies provide pre-existing recognition sites through which the host HIF network can extend oxygen-responsive regulation into the repetitive genome. This mechanism parallels other examples in which TE-derived regulatory sequences have been recruited into host developmental and immune programmes, including ERV-derived enhancers controlling placental and interferon-responsive genes. Rather than representing isolated evolutionary accidents, these cases support a broader principle whereby the regulatory features that once promoted retroelement propagation can subsequently be repurposed to expand host gene-regulatory networks.

The CRISPR experiments extend HIF1A occupancy from biochemical association to regulatory function. Deletion of two independently selected HIF1A-bound LTR7/HERVH loci produced transcriptional consequences, with TG1 in particular affecting a broad local genomic interval. The different magnitude of the TG1 and TG2 responses also argues against treating all HERVH copies as equivalent regulatory units; sequence recognition by HIF1A identifies responsive elements, but their downstream output is likely determined by the genomic environment into which each element inserted. These experiments do not yet identify the promoters contacted directly by TG1 or TG2, nor do they establish how these regulatory effects influence cellular physiology during hypoxia. Resolving these relationships will require locus-resolved chromatin-conformation mapping together with physiological perturbation studies.

### The transition from transformed cancer cell lines to primary immune cells reveals context-dependent regulation

The above findings may also be relevant to the tumour microenvironment, where persistent oxygen limitation coexists with profound changes in immune signalling and TE regulation. The strong hypoxic TE activation observed across several cancer cell lines raises the possibility that oxygen availability contributes to the altered retrotranscriptome of tumour cells^55–57^. Because ERV expression can influence innate immune sensing and tumour immunogenicity, it will be important to determine whether hypoxia-induced TE programmes modify tumour–immune interactions or responses to immunotherapy. More broadly, oxygen availability may provide an environmental input into TE regulation across very different physiological settings, for instance, immune cells entering oxygen-restricted tissues to tumour cells residing within chronically hypoxic microenvironments.

The identity of the hypoxia-responsive TE repertoire was strongly context dependent. LTR7/HERVH was most evident in transformed cell lines, whereas primary blood- and tonsil-derived immune cells retained the broader directional TE response without significant family-level LTR7/HERVH induction. This difference may partly reflect sequencing depth, but it is also consistent with a model in which HIF1A responsiveness is constrained by pre-existing chromatin state and cellular identity.^58–60^. Because TE accessibility is highly lineage specific, the same oxygen-sensing pathway could recruit different repetitive elements in different cellular contexts. Testing this model will require matched HIF1A occupancy and chromatin-accessibility measurements in primary immune populations.

### THE1B and NF-κB: a parallel TE-mediated immune activation circuit

THE1B provides one example of how this context dependence may arise. THE1B is a long terminal repeat of the MaLR group. The THE1 family entered the primate genome more than 40 million years ago, and THE1B itself is restricted to anthropoid primates^61,62^. Most copies survive as solitary LTRs that have kept their promoter and enhancer sequence. These MaLR-derived LTRs retain regulatory activity and have previously been co-opted at loci such as CSF1R^63^ and CRH^62^. In our primary immune-cell data, THE1B induction was strongest in activated tonsil cells, and published CD4⁺ T-cell ChIP-seq places NF-κB over these elements. Together with the enrichment of immune transcription factors at HIF1A-bound regions, this observation suggests that oxygen sensing can intersect with activation-dependent TF networks at repetitive elements. We therefore view HIF1A–LTR7/HERVH not as an isolated switch but as one member of a broader class of environmentally responsive TF–TE interactions.

Our data show that THE1B is significantly induced under hypoxia, specifically in activated tonsil T cells, the one condition combining TCR-driven chromatin opening with hypoxia, and that STAT3 and STAT1 are co-upregulated in the same condition, consistent with the NFκB-THE1B axis observed from cell-line data. THE1B shows the opposite direction in monocytes under hypoxia, but it is upregulated when roxadustat is used, which points to stimulus- and cell-type-specific directional control that is not captured by a simple HIF1A activation model. This stimulus- and cell-type-specific directional control is not captured by a simple HIF1A activation model and instead suggests that the directionality of THE1B depends on whether NF-κB or other competing regulators are dominant at these loci in each cell type. THE1B induction is therefore a context-dependent outcome rather than a general feature of the hypoxic TE response, reinforcing that the hypoxic TE response is not a fixed programme but a set of context-specific outcomes shaped by chromatin state, activation history, and signalling environment. The limited overlap between hypoxia- and interferon-responsive TEs further supports the idea that distinct environmental signals recruit different components of the repetitive genome, even though both perturbations bias TE expression toward activation.

Previous work has established that retroelements can donate regulatory sequences to developmental and immune gene networks; our findings extend this principle to oxygen sensing, showing that an ancient retroviral family has become accessible to one of the most fundamental environmental-response pathways in mammalian physiology. Instead of a single HIF1A–HERVH pathway, the resulting picture is of the repetitive genome as a distributed regulatory substrate through which oxygen, inflammatory signals and cellular identity can be integrated.

### Limitations and open questions

Our ChIP-seq data come from four cancer cell lines, and it is not known whether the same HIF1A binding sites at LTR7/HERVH are accessible and occupied in primary immune cells under hypoxia. Future experiments combining ATAC-seq or ChIP-seq in primary cells with hypoxia treatment would address this directly. An important next step will be to determine whether the HIF1A–LTR7/HERVH interactions identified here operate in physiological hypoxic niches in vivo and which host genes they contact directly. Resolving those relationships will establish whether retroviral sequence co-option has merely expanded the transcriptional repertoire available to HIF1A or has generated lineage-specific physiological functions that cannot be achieved through conventional HIF targets alone. The CRISPR deletion experiments establish that individual TE loci influence nearby gene expression, but the mechanism, whether through enhancer activity, promoter activity, non-coding RNA production, or effects on chromatin topology, is not resolved here. The multi-mapping and drop-out problems from single-cell data for highly repetitive TE families confer a limitation, but the HERVH is quite mappable at the locus-level from bulk-sequencing datasets. Long-read sequencing, which has more ability to assign reads to individual locus copies, would provide a more accurate picture of which specific HERVH loci respond to hypoxia, especially to map the precise transcript structures.

## CONCLUSION

Our findings extend the canonical response to oxygen into the repetitive genome. Hypoxia consistently shifts TE expression toward activation, in striking contrast to the more bidirectional and cell-state-dependent response of host genes. Within this broader retrotranscriptomic response, HIF1A selectively engages HRE-containing LTR7/HERVH loci, and loss of HIF1A preferentially affects the HERVH elements it occupies. CRISPR perturbation further shows that individual HIF1A-bound LTR7/HERVH loci have regulatory consequences for host transcription, although their direct target genes and physiological functions remain to be resolved. These findings identify endogenous retroviral sequences as a regulatory substrate for oxygen sensing and provide an example of molecular exaptation in which regulatory features inherited from ancient retroviral infection have become incorporated into a fundamental mammalian environmental-response network. More broadly, they position the repetitive genome as an interface through which oxygen availability, cellular state and immune signalling can be integrated into human gene regulation.

## METHODS

### Genome index preparation for STAR alignment

To enable simultaneous mapping of gene and TE reads from RNA-seq data, we constructed a custom human genome index integrating RepeatMasker TE annotations with the GENCODE gene reference. The analyses were performed using STAR (v2.7), BEDTools (v2.31), and GffRead (v0.12).

GENCODE human release 49 and the corresponding genome assembly (GRCh38 primary assembly) were downloaded from the GENCODE repository. The annotation was filtered to retain only primary chromosomes (1-22, X, Y, M) and to exclude entries corresponding to rRNA, rRNA pseudogenes, and transcripts labeled with decay-related biotypes. The filtered GTF was converted to BED format to extract exon coordinates for use in subsequent intersection analyses.

To annotate TEs, the hg38 RepeatMasker dataset was parsed to remove low-complexity, simple-repeat, satellite, and RNA gene-related elements, as well as entries shorter than 100 bp. TEs overlapping annotated gene exons were excluded using bedtools intersect to avoid redundancy. The retained elements were reformatted into GTF entries with attributes specifying repName, repClass, and repFamily, and prefixed with the identifier "TE_". The final combined reference GTF, containing both gene and TE annotations, was used to build the STAR genome index.

### RNA-seq alignment and quantification

RNA-seq reads (PRJNA282396, PRJNA606047, PRJNA773654) from cancer cell lines (A549, H460, HeLa, RKO) and HIF1A knockout (HIF1A-KO) cells were aligned to the custom hg38 combined reference genome using STAR (v2.7) in two-pass alignment mode. Alignment parameters were set to allow a maximum of 100 multimapping locations (--outFilterMultimapNmax 100). Read counts at gene and TE loci were extracted using featureCounts (Subread v2.0) with strand-aware counting and multi-overlap assignment enabled. For A549, two independent biological replicates from separate GEO projects (PRJNA606047 and PRJNA773654) were processed independently and merged post-hoc by averaging log2 fold changes and taking the minimum adjusted p-value across the two analyses.

For primary immune cells bulk RNA-seq from blood and tonsil donors, transcript-level quantification was performed using Salmon (v1.9) in mapping-based mode against a decoy-aware index built from the combined gene-and-TE reference. Salmon was run with default parameters with –validateMappings flag enabled. Gene and TE expression values were imported into R using tximeta and summarized at the gene level using tximport with length-scaled TPM.

### Differential expression analysis

Differential expression between hypoxia and normoxia conditions was assessed using DESeq2 (v1.38). For cancer cell line RNA-seq, each cell line was analyzed independently with a simple two-group design (∼condition). For primary CD4+ T cells from blood and tonsil, independent analyses were run for each tissue (blood or tonsil). Multiple testing correction was applied using the Benjamini-Hochberg procedure, with independent hypothesis weighting (IHW) applied where sample sizes permitted. Unless otherwise stated, significance was defined as adjusted p-value < 0.05 with no log2 fold change cutoff for primary analyses. For heatmap feature selection, a stricter threshold of adjusted p-value < 0.01 and |log2FC| > 1 was applied, followed by a consistency filter requiring all samples within at least one condition to agree in the direction of expression deviation from the cross-sample mean.

For CRISPR KO experiments at the TG1 (chr19:15,828,728-15,829,215; hg38) and TG2 chr5:147,869,835-147,870,673; hg38), differential expression between KO and control cells was assessed using DESeq2 with a two-group design. Significance was defined as adjusted p-value < 0.05 and |log2FC| >= 0.5 unless otherwise stated.

### Sign test for directional bias

To assess whether the TE transcriptional response to hypoxia was directionally biased beyond chance, a one-sided binomial sign test was applied to the log2 fold changes of all tested TE families in each comparison, regardless of individual significance. For each comparison, the number of TE families with log2FC > 0 was tested against a null expectation of 50%.

### Gene ontology enrichment analysis

Gene ontology over-representation analysis was performed using clusterProfiler (v4.6) with the org.Hs.eg.db annotation database. Input gene sets consisted of significantly upregulated or downregulated genes (adjusted p-value < 0.05) from each comparison. GO terms from Biological Process (BP), Molecular Function (MF), and Cellular Component (CC) ontologies were tested. Results were filtered for adjusted p-value < 0.05 and redundant terms were collapsed using the simplify function with a semantic similarity cutoff of 0.7 (Wang measure). Gene ratio was defined as the proportion of input genes annotated to each GO term relative to the total number of input genes with GO annotation.

### ChIP-seq data processing

ChIP-seq data for HIF1A across multiple cell lines (A549, H460, HCT116, and RKO) under normoxic and hypoxic conditions were obtained from publicly available datasets (PRJNA606242). Raw sequencing reads were aligned to the human reference genome hg38 using Bowtie2 in very-sensitive-local mode. Peak summits were derived from processed peak files and unified across samples using bedtools merge (200 bp merging distance) to generate a non-redundant catalogue of HIF1A binding sites. All analyses were restricted to standard chromosomes (chr1-22, X, Y), and summit positions were extended ±50 bp for motif analysis.

In addition to the HIF1A-focused analysis, a separate transcription factor enrichment analysis was performed using publicly available ENCODE ChIP-seq peak datasets. These data were processed independently and peak summits were extracted and aggregated per TF for downstream enrichment analysis. TF datasets were curated to exclude replication, repair, and basal transcription machinery factors, retaining 188 biologically informative TF datasets. TF classification was obtained by parsing the TFClass database hierarchy.

### HIF1A ChIP-seq peak annotation and TE overlap analysis

Peaks were annotated with respect to genomic features (promoter ±1 kb, gene body, intergenic) using ChIPseeker (v1.34). TE loci overlapping HIF1A peaks were extracted and their family, class, and genomic context recorded. The genome-wide background for enrichment comparisons was the filtered RepeatMasker GTF used for RNA-seq quantification, restricted to TEs longer than 100 bp and not overlapping annotated gene exons.

### Permutation-based TE enrichment analysis

Enrichment of HIF1A ChIP-seq peaks at each TE family was assessed using the regioneR package (v1.44) with 1,000 permutations. For each TE family, the observed overlap between HIF1A summit positions and TE loci was compared against a null distribution generated by randomly repositioning HIF1A peaks across the genome while preserving chromosome-level peak density. Enrichment p-values were computed from the permutation null distribution.

### Motif enrichment analysis

Sequence motif enrichment within HIF1A-bound LTR7 loci was assessed using MEME-SEA (v5.5.9). Sequences of ±50 bp around HIF1A ChIP-seq summit positions overlapping LTR7 loci were used as the positive set, and sequences centered on LTR7 loci with no evidence of HIF1A binding were used as the negative set. The JASPAR 2026 vertebrate motif database was used as the reference motif collection. Enrichment was assessed using a one-sided Fisher’s exact test as implemented in MEME-SEA, with E-value correction for multiple testing. Fold enrichment was defined as the ratio of the fraction of positive sequences containing the motif to the fraction of negative sequences containing the motif.

### TF co-localization enrichment analysis

Co-localization enrichment between HIF1A ChIP-seq summit positions and 188 curated ENCODE transcription factor ChIP-seq datasets was assessed analytically. For each TF, the observed number of TF summit positions falling within ±10 bp of a HIF1A summit was counted. The expected overlap under independence was computed analytically from the fraction of each chromosome’s length covered by HIF1A summit windows, multiplied by the number of TF summits on that chromosome, and summed across chromosomes. Fold enrichment was defined as observed divided by expected overlap. Statistical significance was assessed using a one-sided binomial test and corrected for multiple testing across 188 TFs using the Benjamini-Hochberg method.

### BD Rhapsody single-cell multiomics

PBMCs from three healthy donors were cultured under four conditions: normoxia (NOX), hypoxia (HOX, 1% O2), pharmacological HIF1A stabilization with roxadustat, and interferon stimulation for 24 hours. Single-cell whole-transcriptome and AbSeq protein sequencing was performed using the BD Rhapsody system following the manufacturer’s protocol. Sample multiplexing was performed using BD Sample Tags, and doublet removal was performed using the BD Rhapsody analysis pipeline. The resulting read data were aligned to a combined gene-and-TE reference. Cell clustering and annotation were performed in Seurat (v5.0.2) using Harmony (v1.2.0) for batch correction across donors. Cell types were annotated using a combination of AbSeq protein markers and canonical RNA lineage markers. Family-level and locus-level TE differential expression was assessed using the bimod likelihood ratio test implemented in FindMarkers from the Seurat package.

### Blood and tonsil experiments

Primary immune cells were isolated from three healthy blood donors and tonsil samples was obtained from four donors. For activation experiments, cells were stimulated with anti-CD3/CD28 coated beads. Hypoxia was induced by culturing cells at 1% O2 in a hypoxic chamber for 24 hours. RNA was extracted and used to prepare RNA-seq libraries, which were then sequenced.

### CRISPR deletion experiments

To functionally assess HIF1A-bound HERVH elements, two HERVH loci overlapping HIF1A ChIP-seq peaks were selected for CRISPR-mediated deletion and are referred to here as TG1 and TG2. hESC-H1 and hESC-H9 cells were transfected with Cas9-gRNA constructs (Addgene plasmid #86986) targeting the respective loci. One day after transfection, single cells were sorted into 96-well plates. Cells were maintained with medium changes every other day and supplemented with CloneR (STEMCELL Technologies) instead of ROCK inhibitor during clonal outgrowth. Colonies were expanded for approximately 3 weeks before transfer to 12-well plates.

Genomic DNA was isolated from individual clones, and deletion of the targeted HERVH locus was assessed by PCR across the edited interval, followed by agarose-gel electrophoresis and Sanger sequencing of candidate knockout clones. Successfully edited clones were expanded for downstream transcriptional analyses. For each targeting experiment, a clone that underwent the same Cas9-gRNA transfection, single-cell sorting and clonal expansion procedure but retained the intact HERVH locus, as confirmed by PCR, was used as a matched control.

### Cis-regulatory and evolutionary conservation analysis

To assess the cis-regulatory footprint of each CRISPR-deleted TE locus, the genomic coordinates of all expressed genes were intersected with the deleted locus coordinates. Genes within 1 Mb windows were extracted and their differential expression values from the corresponding KO experiment were plotted against their distance from the deleted locus midpoint.

## Data availability

Cancer cell line RNA-seq and HIF1A ChIP-seq data were downloaded from the Gene Expression Omnibus (GEO) from the following accession numbers: PRJNA606047, PRJNA773654, PRJNA282396, and PRJNA606242. Primary blood- and tonsil-derived immune cell RNA-seq and BD Rhapsody single-cell multiomics data generated in this study have been deposited in the Gene Expression Omnibus and will be released upon publication. Accession numbers will also be listed at https://github.com/umutcakir/Hypoxia_HIF1A_targets. The supporting tables (Supplementary Tables S1-S9) and Figures (Supplementary Figures S1-S11) are available at the Zenodo (zenodo.org/records/22164490; DOI: doi.org/10.5281/zenodo.22164490).

## Code availability

All custom scripts used for data processing, statistical analysis, and figure generation in this study are available at https://github.com/umutcakir/Hypoxia_HIF1A_targets.

## Acknowledgements

This work was supported by the Centre National de la Recherche Scientifique (CNRS) through a Chaire de Professeur Junior (CPJ) awarded to M.S. for the project “Biologie des systèmes en physiopathologie.” and FRNEM welcome grant (FRN202509051056). UC received support from the IMPRS-Genome Science PhD program. H.E. is supported by the Deutsche Forschungsgemeinschaft (DFG, German Research Foundation) SFB TRR 274/2 - 408885537 project C01.

## Author Contributions

Concept, design, supervision: MS, RS; Funding acquisition: MS, RS, ZI; Drafting manuscript and display items: MS, UC; Data acquisition and generation: UC, SYA, JF, SW, HS, RS, FM, ZI, MS; Data analyses & interpretation: UC, SYA, FM, HE.

All authors read and approved the final version of the manuscript.

## Figure legends

**Figure S1.**
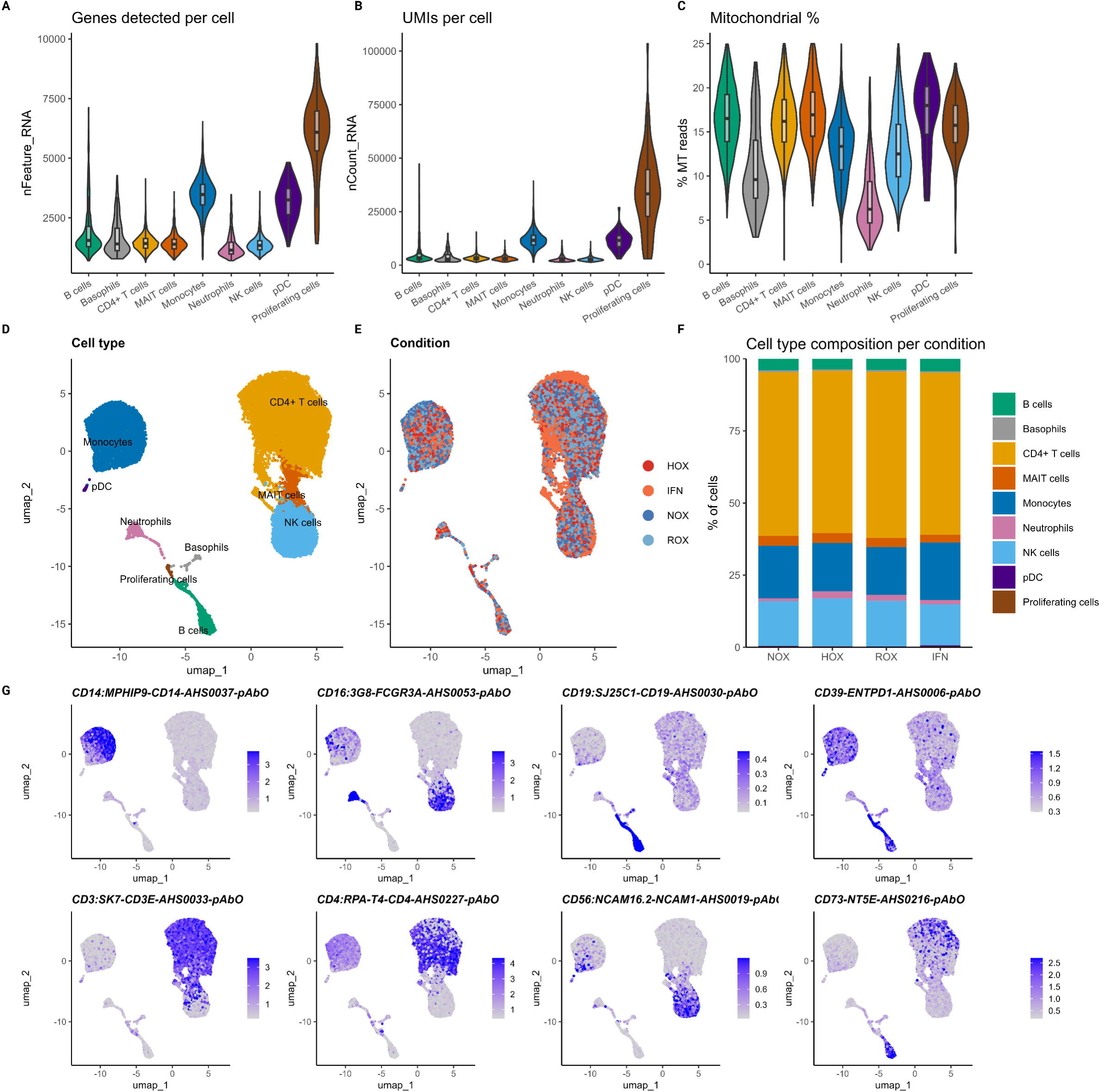
PBMC single-cell quality control, cell type annotation, and AbSeq protein expression. **(A)** Genes detected per cell by annotated cell type. **(B)** Unique molecular identifiers per cell by cell type. **(C)** The percentage of mitochondrial reads per cell by cell type. **(D)** UMAP coloured by annotated cell type. **(E)** UMAP coloured by experimental condition. **(F)** Cell type composition per condition. **(G)** AbSeq surface protein expression projected onto the UMAP for the markers used in annotation: CD14, CD16, CD19, CD39, CD3, CD4, CD56, and CD73.

**Figure S2.**
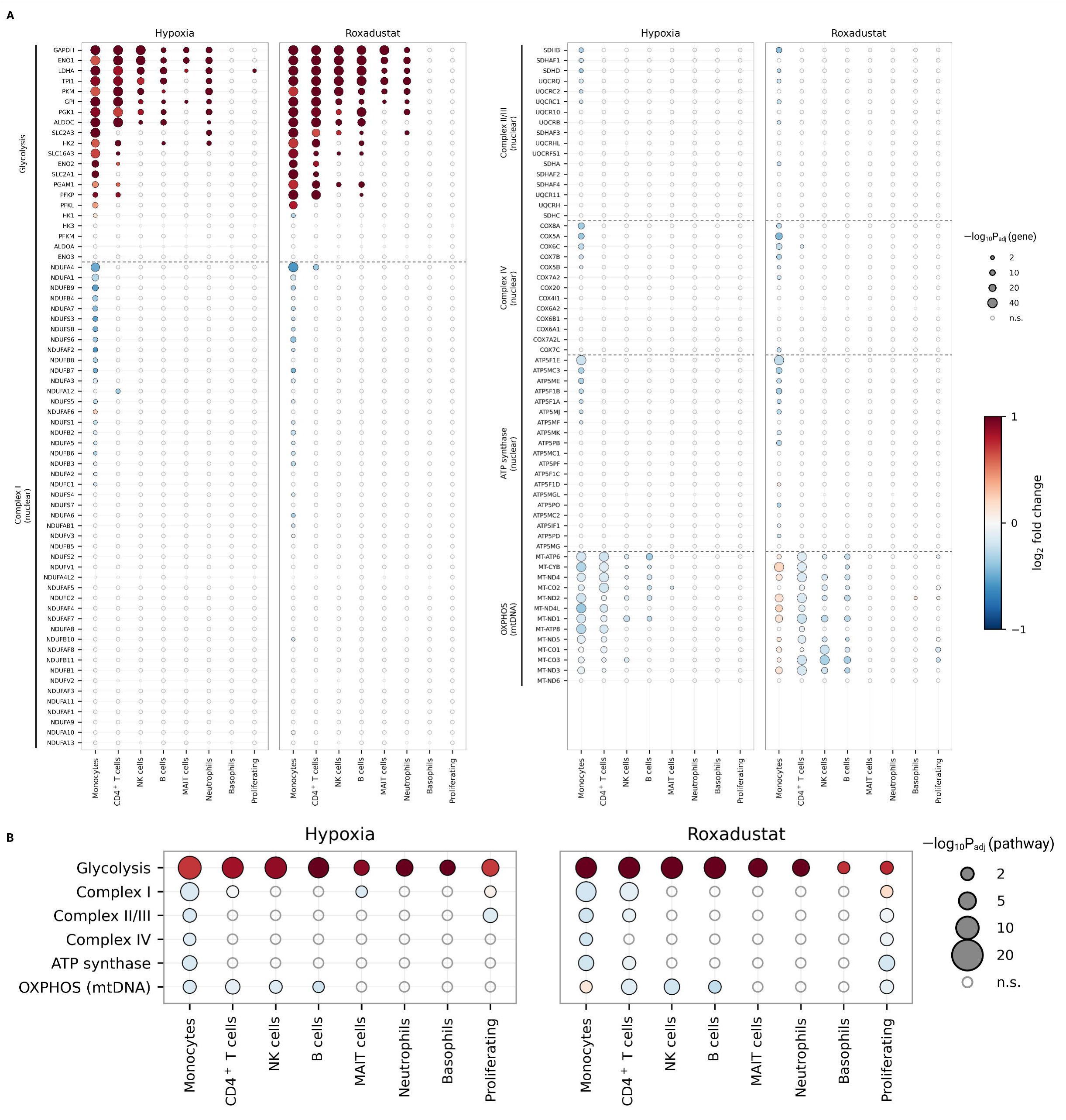
Gene-level and pathway-level metabolic responses to hypoxia and roxadustat across PBMC populations. **(A)** Log₂ fold change for individual glycolytic genes and for nuclear- and mitochondrially encoded OXPHOS subunits across eight annotated populations, under hypoxia which is shown on the left and roxadustat which is shown on the right. Point size reflects −log₁₀ padj. **(B)** Pathway-level summary of the (A) data. Each pathway is tested by comparing the log₂ fold changes of all its tested genes against the distribution of all tested genes in the same cells. Glycolysis is significantly shifted upward across all populations under both stimuli, whereas the respiratory complexes are, in general, shifted downward.

**Figure S3.**
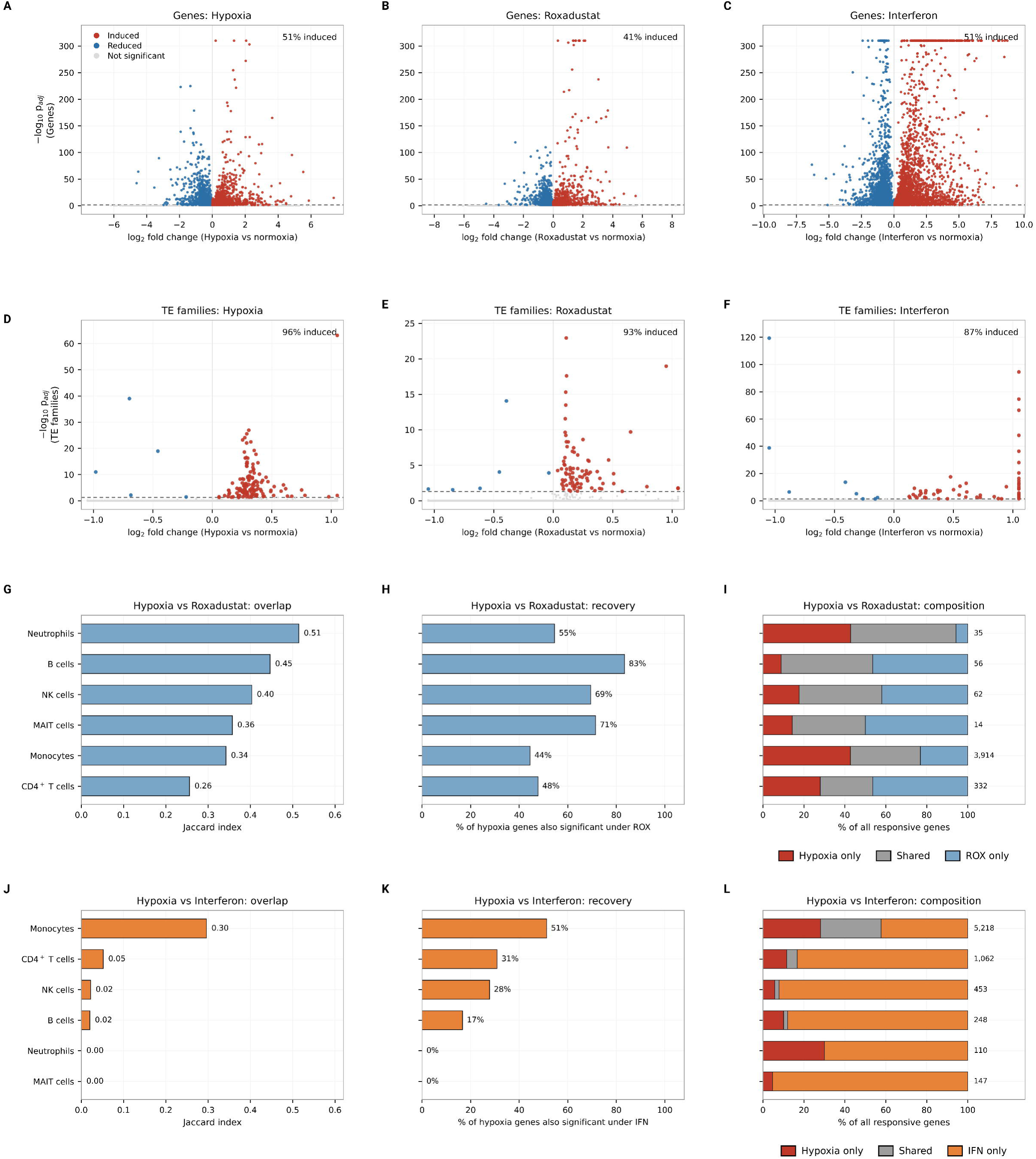
Differential expression and stimulus comparisons in single-cell PBMC data. **(A)** Volcano plot of gene-level differential expression under hypoxia versus normoxia, with the percentage of significant genes induced given at the top right. **(B)** Volcano plot of gene-level for roxadustat versus normoxia. **(C)** Volcano plot of gene-level for interferon versus normoxia. **(D)** Volcano plot of TE family differential expression under hypoxia versus normoxia. **(E)** Volcano plot of TE family for roxadustat versus normoxia. **(F)** Volcano plot of TE family for interferon versus normoxia. TE responses are induced by 87% to 96% across the three stimuli, while gene responses remain approximately balanced. **(G)** Jaccard overlap between the hypoxia- and roxadustat-responsive gene sets per cell type. **(H)** The percentage of hypoxia-responsive genes is also significant under roxadustat. **(I)** Composition of all responsive genes per cell type, as hypoxia only, shared, or roxadustat only, with the total per cell type is shown on the right. **(J)** Jaccard overlap between the hypoxia- and interferon-responsive gene sets per cell type. **(K)** The percentage of hypoxia-responsive genes is also significant under interferon. **(L)** Composition of all responsive genes per cell type, as hypoxia only, shared, or interferon only, with the total per cell type is shown on the right.

**Figure S4.**
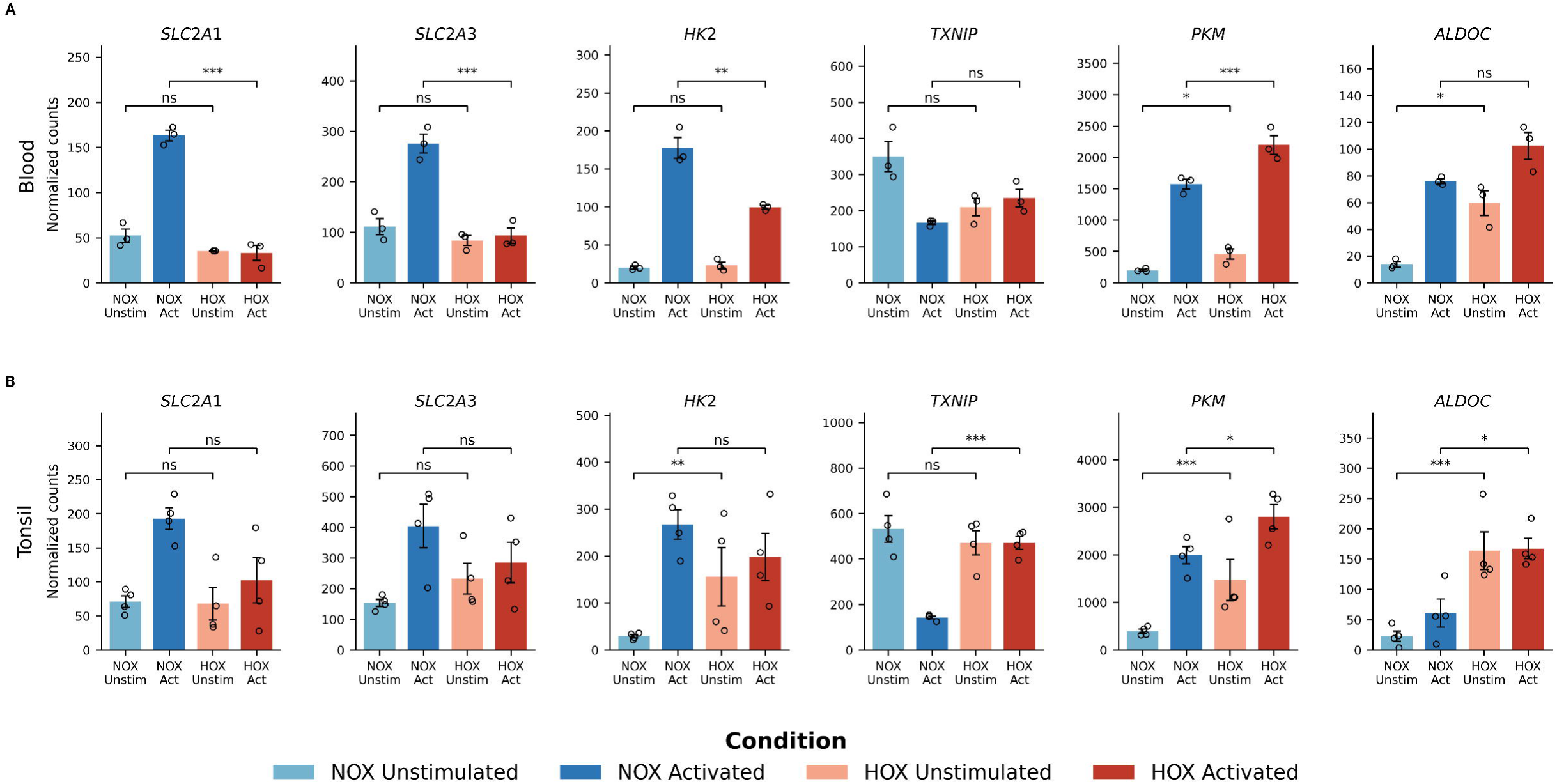
Metabolic gene expression across hypoxia and activation conditions in blood and tonsil derived immune cells. **(A)** Normalised counts for SLC2A1, SLC2A3, HK2, TXNIP, PKM, and ALDOC across four conditions in the blood-derived cells. Points are individual donors, and error bars show standard error. Brackets indicate the comparison between hypoxia and normoxia within each stimulation state. **(B)** The same plots for the tonsil-derived cells. SLC2A1, SLC2A3, and HK2 are reduced by hypoxia only after activation, whereas TXNIP retains its resting level under hypoxia rather than declining as it does under normoxia.

**Figure S5.**
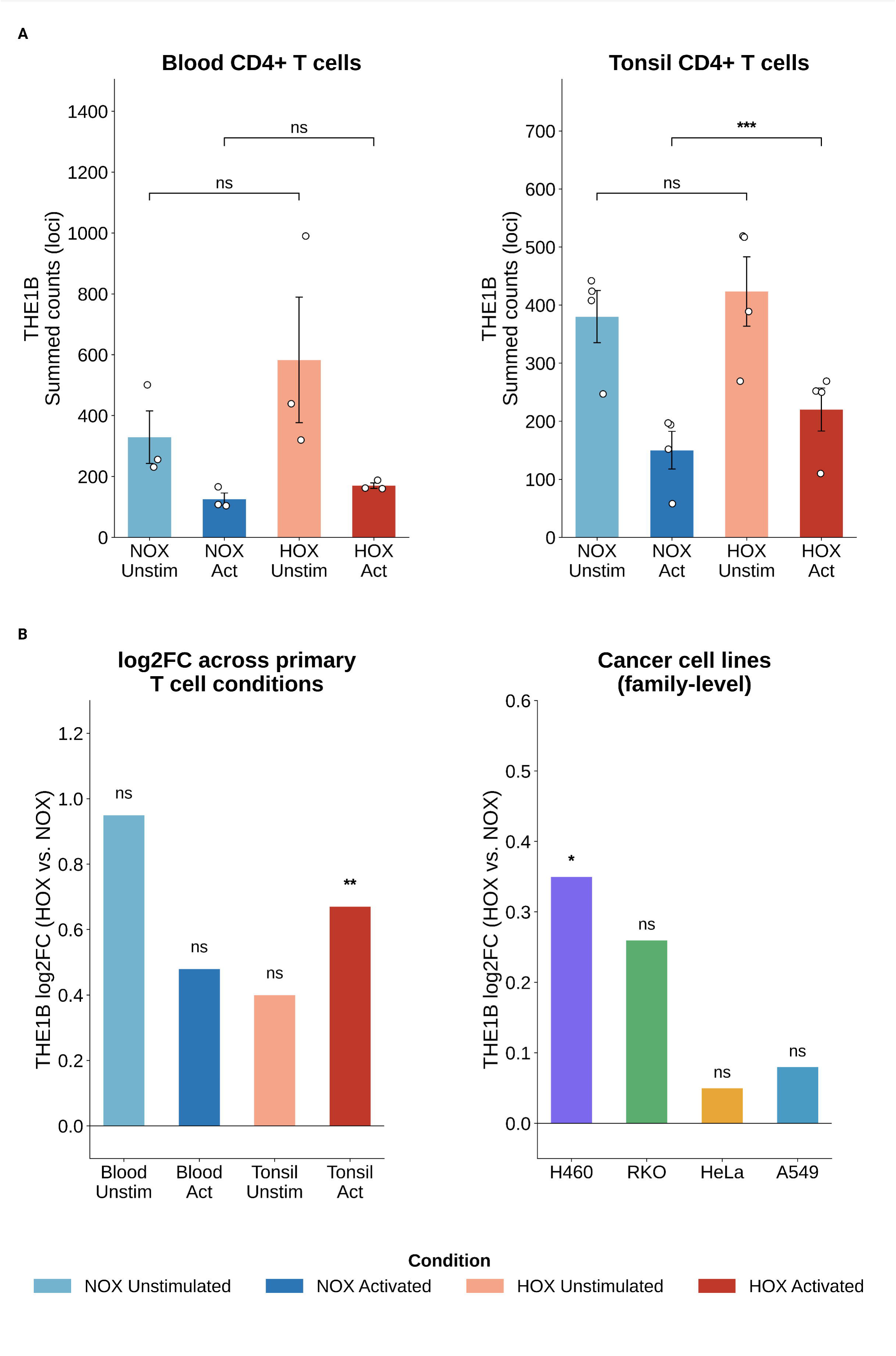
THE1B expression in primary blood and tonsil-derived cells and cancer cell lines. **(A)** Summed locus-level THE1B counts across four conditions in blood, which is shown on the left and tonsil shown on the right. Points are individual donors and error bars show standard error. **(B)** THE1B log₂ fold change (hypoxia versus normoxia) across the four primary cell comparisons (left) and the four cancer cell lines, shown on the left and right, respectively. THE1B is positive in every comparison and reaches significance in the activated tonsil and in H460 cell line.

**Figure S6.**
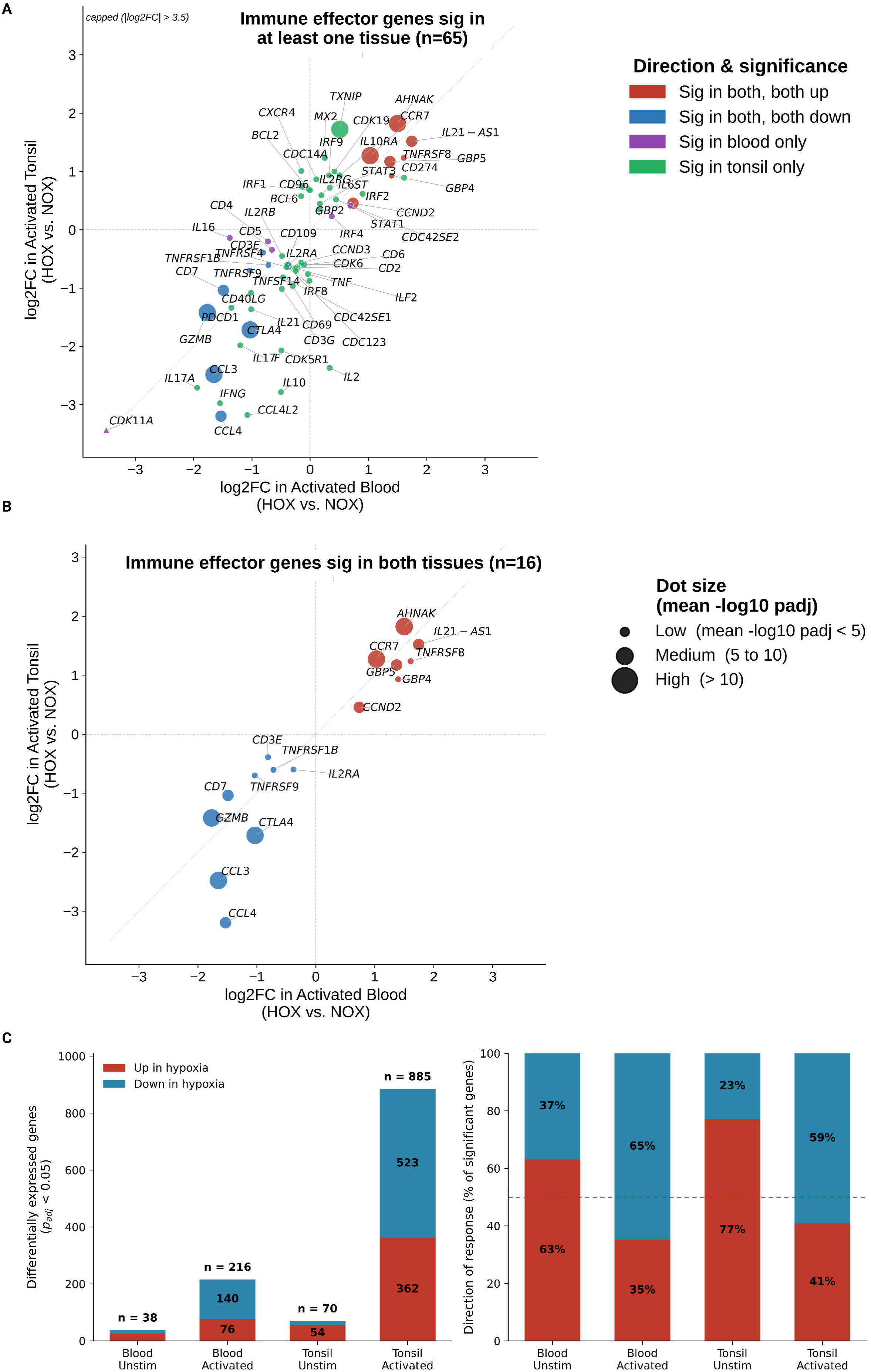
Immune effector gene responses to hypoxia in activated blood and tonsil derived immune cells. **(A)** Log₂ fold change under hypoxia in activated blood against activated tonsil for immune effector genes significant in at least one tissue. The colour code indicates direction and in which tissue significance was reached. **(B)** The same plot restricted to the 16 genes that are significant in both tissues. Dot size reflects the mean −log₁₀ padj across the two tissues. All 16 agree on the direction between tissues. **(C)** Number of significantly differentially expressed genes per comparison and the direction of the response as a percentage of significant genes, which are shown on the left and right, respectively.

**Figure S7.**
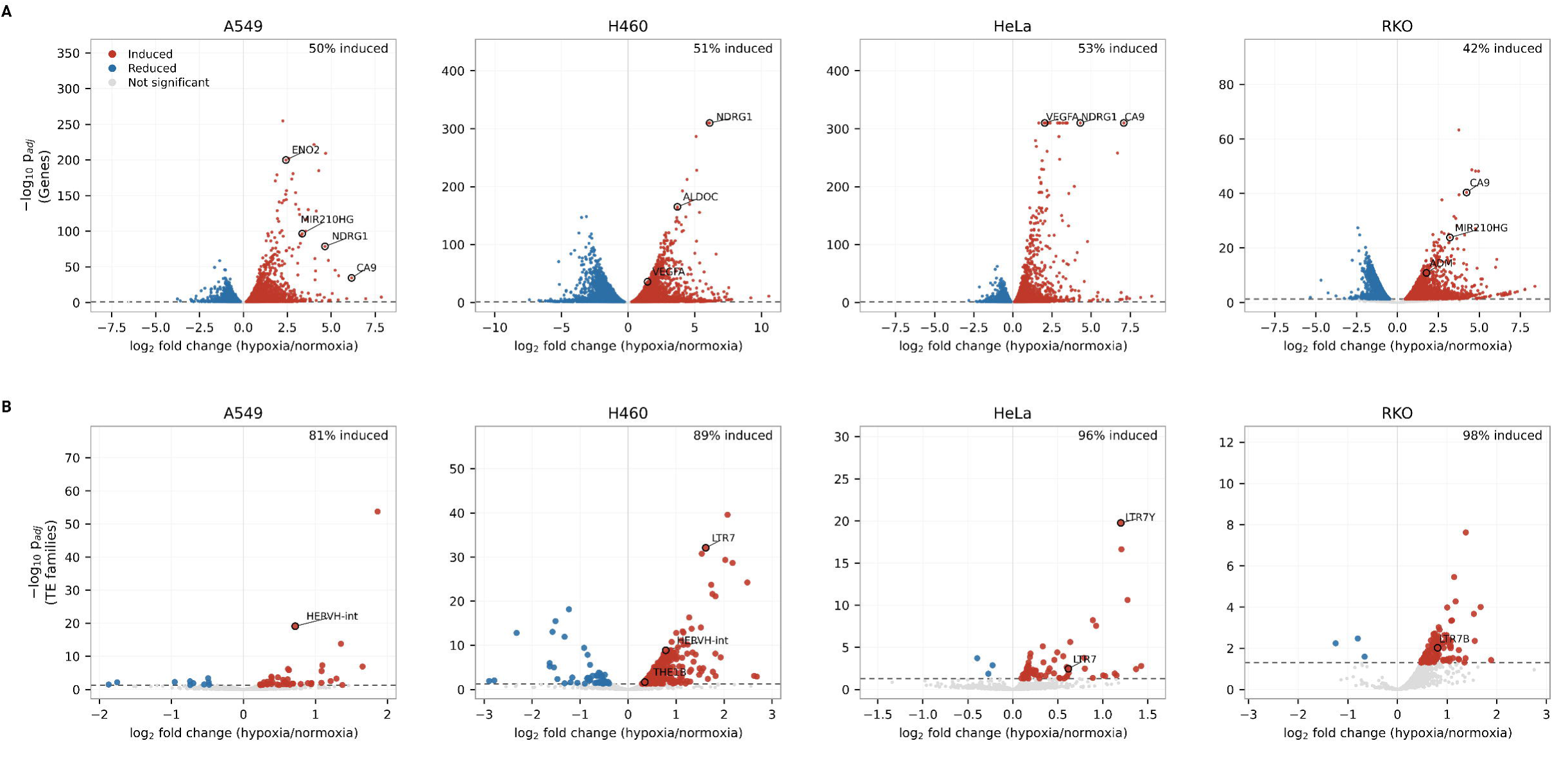
Differential expression in cancer cell lines under hypoxia. **(A)** Volcano plots of gene-level differential expression (hypoxia versus normoxia) in A549, H460, HeLa, and RKO cells, with the percentage of significant genes induced given at the top right of each panel. Selected canonical hypoxia related genes are labelled. **(B)** Volcano plots of TE family differential expression in the same cell lines. The percentage induced ranges from 81% to 98%. Selected TE families are labelled.

**Figure S8.**
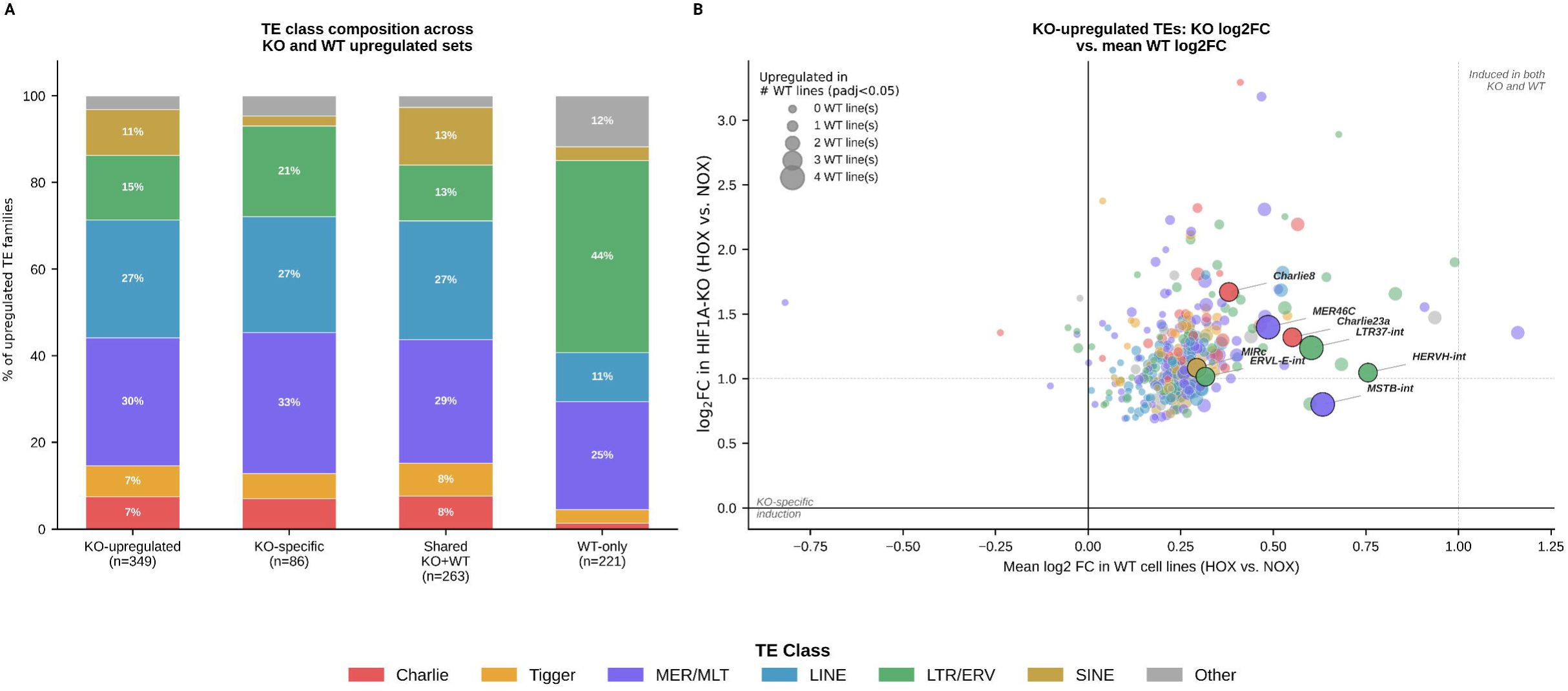
TE class composition and hypoxia-responsiveness of TE families were upregulated in HIF1A knockout cells. **(A)** Class composition of upregulated TE features across four sets: all families upregulated in the HIF1A knockout, those specific to the knockout, those shared between the knockout and at least one wild-type cell line, and those upregulated in wild-type lines only. Group sizes are given below each bar. **(B)** Log₂ fold change in the HIF1A knockout against the mean log₂ fold change across wild-type cell lines for each knockout-upregulated TE family. Point size reflects the number of wild-type lines in which the family is significantly upregulated, and colour indicates the TE class. Selected families are labelled.

**Figure S9.**
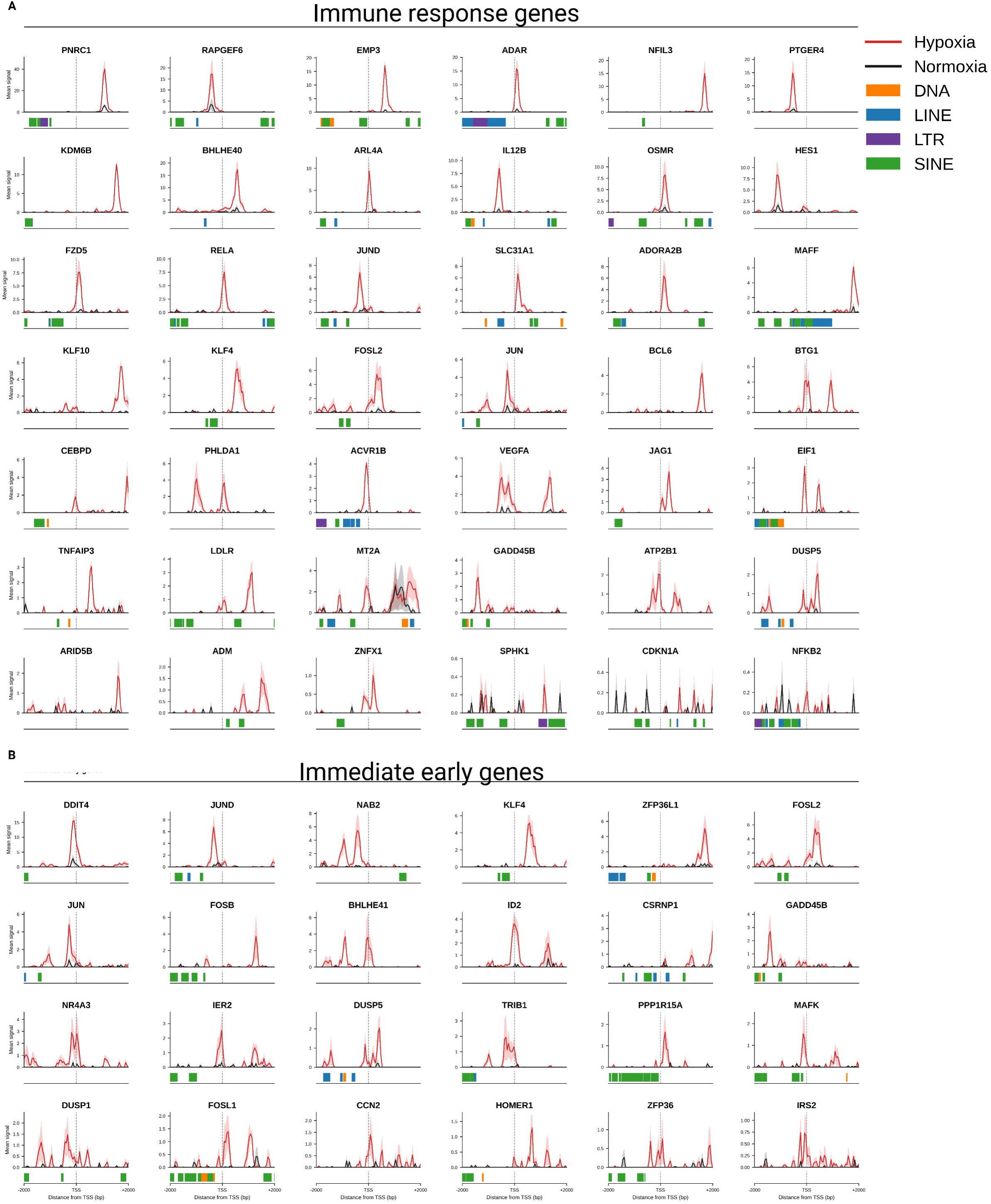
HIF1A ChIP-seq signal at gene promoters, averaged across cell lines. **(A)** Mean ChIP-seq signal in a ±2 kb window around the transcription start site of each of selected 42 immune response genes under hypoxia (red) and normoxia (black), averaged across the four cell lines. Shaded bands show the range between lines. TE loci within each window are annotated by class, which are shown below the x-axis. **(B)** The same plot but for the selected 24 immediate early genes.

**Figure S10.**
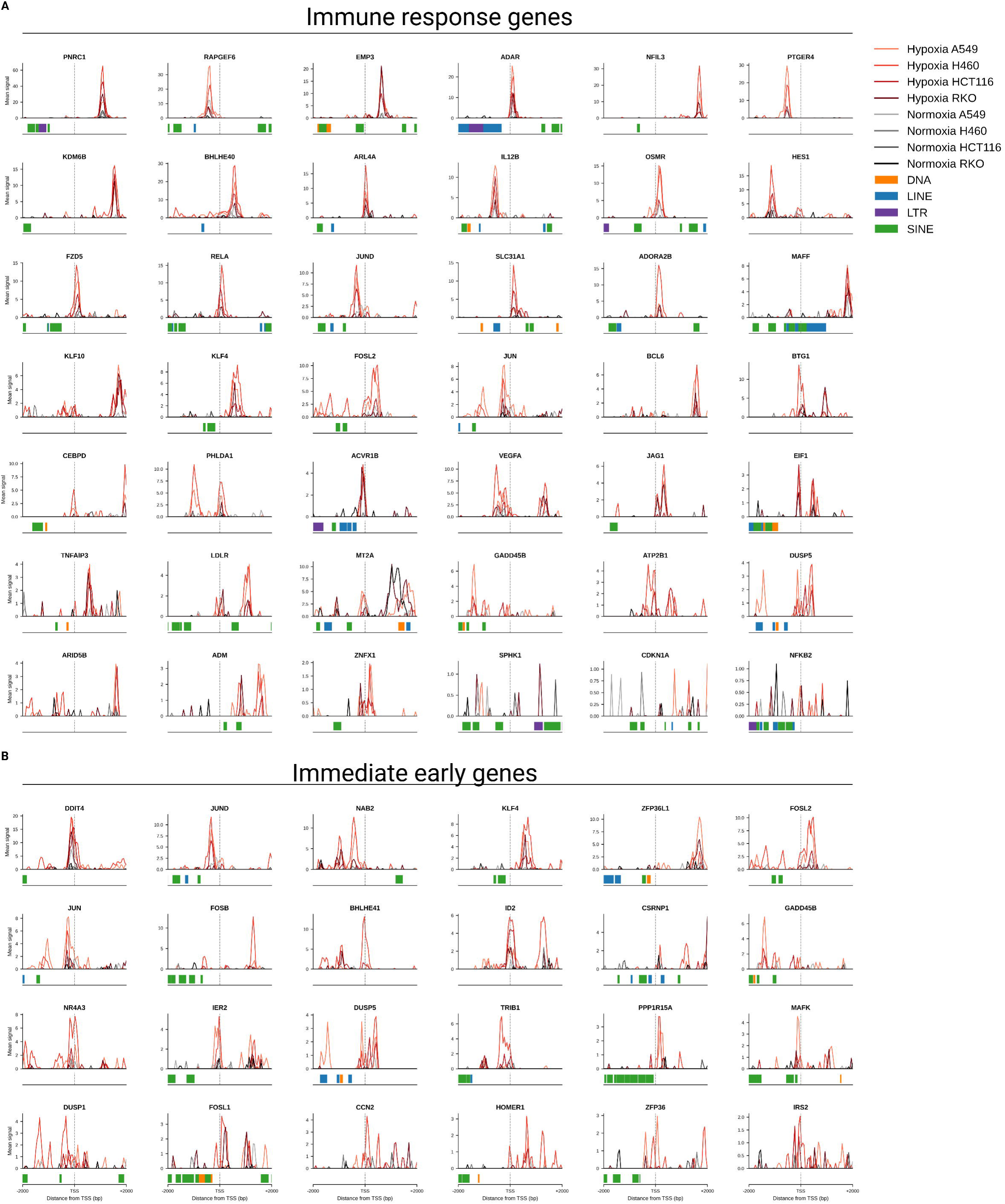
HIF1A ChIP-seq signal at gene promoters, resolved by cell line. **(A)** Mean ChIP-seq signal in a ±2 kb window around the transcription start site of each of 42 immune response genes, shown separately for A549, H460, HCT116 and RKO under hypoxia and normoxia. TE loci within each window are annotated by class below the x-axis. **(B)** The same for the 24 immediate early genes.

**Figure S11.**
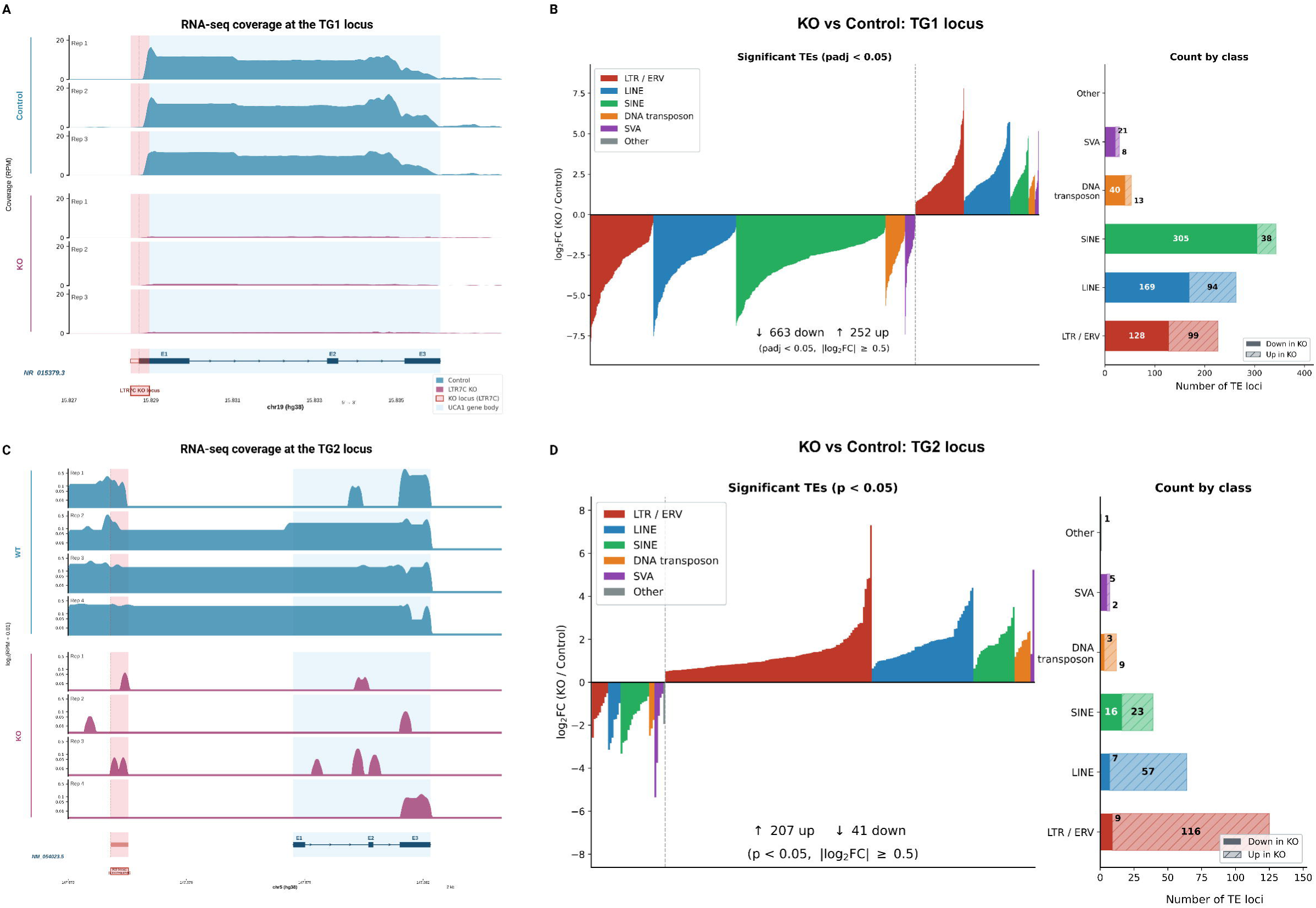
RNA-seq coverage and TE-level responses at the TG1 and TG2 loci. **(A)** RNA-seq coverage across the TG1 locus in control and knockout replicates. The deleted element is shaded in pale red and the closest gene to TG1 locus is shown in blue. **(B)** Left: waterfall plot of all significantly differentially expressed TE loci at the TG1 locus, ranked by log₂ fold change and coloured by TE class. The right bar plot shows the counts by class and direction. **(C)** RNA-seq coverage across the TG2 locus in control and LTR7/HERVH knockout replicates, plotted in the same way as in (A). **(D)** Waterfall plot and class counts for the TG2 locus.

## Supplementary Tables

Supplementary Table S1. Gene and TE family differential expression results from bimod test across cell types in scRNA-seq PBMC analysis and pairwise comparisons (HOX vs. NOX, ROX vs. NOX, IFN vs. NOX, ROX vs. HOX).

Supplementary Table S2. DESeq2 expression results (HOX vs. NOX) for genes and TE families in primary immune cells across four comparisons: blood unstimulated, blood activated, tonsil unstimulated, and tonsil activated, together with a gene by sample TPM matrix for all sequenced samples.

Supplementary Table S3. DESeq2 differential expression results for genes and TE families in each cancer cell line (RKO, A549, H460, HeLa) and HIF1A-KO cells under hypoxia versus normoxia. The sample metadata sheet shows accession numbers and experimental conditions.

Supplementary Table S4. Catalog of HIF1A ChIP-seq peaks overlapping TE loci. They are annotated with peak coordinates, TE family, TE class, and TE name.

Supplementary Table S5. Permutation-based enrichment test results (regioneR, 1,000 permutations) for HIF1A ChIP-seq peak overlap at each TE family, reported separately for each cell line and the combined dataset.

Supplementary Table S6. MEME-SEA motif enrichment results comparing HIF1A-bound LTR7 sequences against unbound LTR7 sequences, including motif identity, enrichment statistics, and the proportion of positive and negative sequences carrying each motif.

Supplementary Table S7. Co-localization enrichment statistics for 188 ENCODE transcription factors at HIF1A ChIP-seq binding sites, including observed and expected overlap counts, fold enrichment, and adjusted p-values.

Supplementary Table S8. DESeq2 results for genes and TE loci in TG1 (KO versus control).

Supplementary Table S9. DESeq2 results for genes and TE loci in TG2 (KO versus control).

